# A unified framework of realistic in silico data generation and statistical model inference for single-cell and spatial omics

**DOI:** 10.1101/2022.09.20.508796

**Authors:** Dongyuan Song, Qingyang Wang, Guanao Yan, Tianyang Liu, Jingyi Jessica Li

## Abstract

In the single-cell and spatial omics field, computational challenges include method benchmarking, data interpretation, and in silico data generation. To address these challenges, we propose an all-in-one statistical simulator, scDesign3, to generate realistic single-cell and spatial omics data, including various cell states, experimental designs, and feature modalities, by learning interpretable parameters from real datasets. Furthermore, using a unified probabilistic model for single-cell and spatial omics data, scDesign3 can infer biologically meaningful parameters, assess the goodness-of-fit of inferred cell clusters, trajectories, and spatial locations, and generate in silico negative and positive controls for benchmarking computational tools.

## Introduction

Single-cell and spatial omics technologies have provided unprecedented multi-modal views of individual cells. As the earliest single-cell technologies, single-cell RNA-seq (scRNA-seq) enabled the measurement of transcriptome-wide gene expression levels and the discovery of novel cell types and continuous cell trajectories [1, 2]. Later, other single-cell omics technologies were developed to measure additional molecular feature modalities, including single-cell chromatin accessibility (e.g., scATAC-seq [3] and sci-ATAC-seq [4]), single-cell DNA methylation [5], and single-cell protein abundance (e.g., single-cell mass cytometry [6]). More recently, single-cell multi-omics technologies were invented to simultaneously measure more than one feature modality, such as SNARE-seq (gene expression and chromatin accessibility) [7] and CITE-seq (gene expression and surface protein abundance) [8]. In parallel to single-cell omics, spatial transcriptomics technologies were advanced to profile gene expression levels with spatial location information of cell neighborhoods (i.e., multi-cell resolution; e.g., 10x Visium [9] and Slide-seq [10]), individual cells (i.e., single-cell resolution; e.g., Slide-seqV2 [11]), or sub-cellular components (i.e., sub-cellular resolution; e.g., MERFISH [12]).

Thousands of computational methods have been developed to analyze single-cell and spatial omics data for various tasks [13], making method benchmarking a pressing challenge for method developers and users. Fair benchmarking relies on comprehensive evaluation metrics that reflect real data analytical goals; however, meaningful metrics usually require ground truths that are rarely available in real data. (For example, most real datasets contain “cell types” obtained by cell clustering and manual annotation without external validation; using such “cell types” as ground truths would biasedly favor the clustering method used in the original study.) Therefore, fair benchmarking demands in silico data that contain ground truths and mimic real data, calling for realistic simulators.

The demand for realistic simulators motivated two recent benchmark studies, in which 12 and 16 scRNA-seq simulators were evaluated [14, 15]. Due to the complexity of scRNA-seq data, these benchmarked simulators all require training on real scRNA-seq data, and they are more realistic than the de novo simulators that use no real data but generate synthetic data from theoretical models [15].

Although the benchmark studies found that the simulators scDesign2 [16], ZINB-WaVE [17], and muscat [18] can generate realistic scRNA-seq data from discrete cell types [14, 15], few simulators can generate realistic scRNA-seq data from continuous cell trajectories by mimicking real data [15, 19–22]. Moreover, realistic simulators are lacking for single-cell omics other than scRNA-seq, not to mention single-cell multi-omics and spatial transcriptomics. (To our knowledge, simATAC is the only scATAC-seq simulator that learns from real data, but it can only generate discrete cell types [23].) Hence, a large gap exists between the diverse benchmarking needs and the limited functionalities of existing simulators.

To fill in the gap, we introduce scDesign3, a realistic and most versatile simulator to date. As Fig. 1a shows, scDesign3 can generate realistic synthetic data from diverse settings, including cell latent structures (discrete cell types and continuous cell trajectories), feature modalities (e.g., gene expression, chromatin accessibility, methylation, protein abundance, and multi-omics), spatial locations, and experimental designs (batches and conditions). Note that the predecessor scDesign2 [16] is a special case of scDesign3 for generating scRNA-seq data from discrete cell types; a detailed comparison of scDesign3 with the previous two versions (scDesign [24] and scDesign2) is in Table S1. To our knowledge, scDesign3 offers the first probabilistic model that unifies the generation and inference for single-cell and spatial omics data. Equipped with interpretable parameters and a model likelihood, scDesign3 is beyond a versatile simulator and has unique advantages for generating customized in silico data, which can serve as negative and positive controls for computational analysis, and for assessing the goodness-of-fit of inferred cell clusters, trajectories, and spatial locations in an unsupervised way (Fig. 2a).

**Figure 1:**
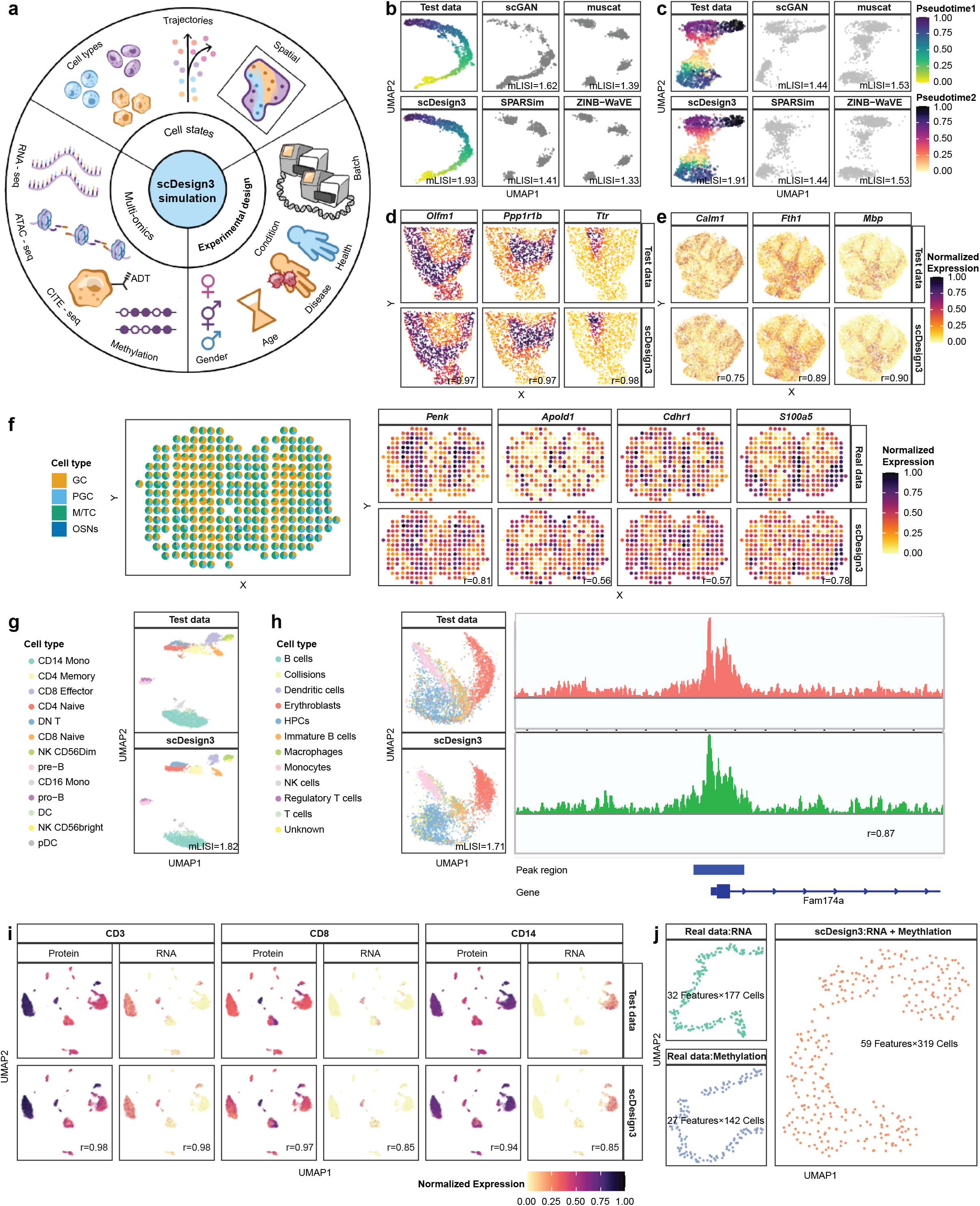
scDesign3 generates realistic synthetic data of diverse single-cell and spatial omics technologies. **a**, An overview of scDesign3’s simulation functionalities: cell states (e.g., discrete types, continuous trajectories, and spatial locations); multi-omics modalities (e.g., RNA-seq, ATAC-seq, and CITE-seq); experimental designs (e.g., batches and conditions). **b–c**, scDesign3 outperforms existing simulators scGAN, muscat, SPARSim, and ZINB-WaVE in simulating scRNA-seq datasets with a single trajectory (b) and bifurcating trajectories (c). Larger mLISI values represent better resemblance between synthetic data and test data. **d–e**, scDesign3 simulates realistic gene expression patterns for spatial transcriptomics technologies 10x Visium (d) and Slide-seq (e). Large Pearson correlation coefficients (*r*) represent similar spatial patterns in synthetic and test data. **f**, using paired scRNA-seq data and spatial transcriptomics data (from the MOB dataset) as input, we define the “ground-truth” cell-type proportions at each spot (left). Each color represents a cell type. With the cell-type proportions, scDesign3 generates synthetic spatial transcriptomics data in which every spot is a mixture of synthetic single cells, given the spot’s cell-type proportions. The four cell-type marker genes exhibit similar spatial expression patterns in real data (right top) and synthetic data (right bottom). Large *r* values represent similar expression patterns in synthetic and test data. **g**, scDesign3 simulates a realistic scATAC-seq dataset at the count level. **h**, scDesign3 simulates a realistic sci-ATAC-seq dataset at both the count level (left: UMAP visualizations of real and synthetic cells in terms of peak counts) and the read level (right: pseudobulk read coverages; coupled with scReadSim [30]). **i**, scDesign3 simulates realistic CITE-seq data. Four genes’ protein and RNA abundances are shown on the cell UMAP embeddings in test data (top) and the synthetic data (bottom). Large *r* values represent similar expression patterns in synthetic and test data. **j**, scDesign3 generates a multi-omics (RNA expression + DNA methylation) dataset (right) by learning from real data that only have a single modality (left). The synthetic data preserve the linear cell topology.

**Figure 2:**
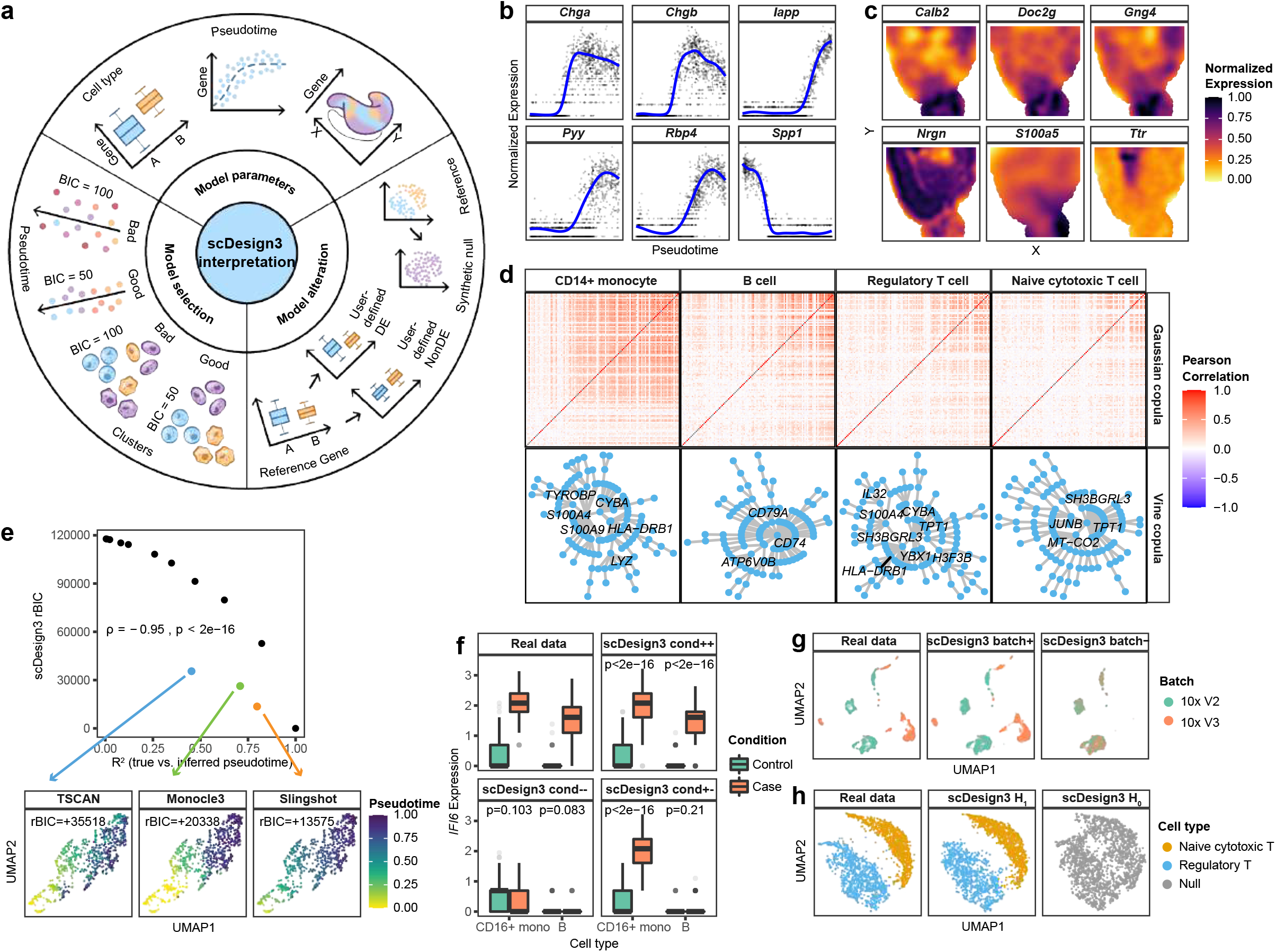
scDesign3 enables comprehensive interpretation of real data. **a**, An overview of scDesign3’s interpretation functionalities based on its model parameters, model selection capacity, and model alteration capacity. **b**, scDesign3 estimates six genes’ expression trends along cell pseudotime that indicates cell differentiation (scRNA-seq dataset PANCREAS in Table S2). **c**, scDesign3 estimates six genes’ expression trends across spatial locations (10x Visium spatial dataset VISIUM in Table S2). **d**, scDesign3 estimates cell-type-specific gene correlations in four cell types (scRNA-seq dataset ZHENGMIX4 in Table S2): pairwise gene correlation matrices by Gaussian copula (top); vine representations by vine copula (bottom), with genes in the first layer (roughly the genes strongly correlated) labeled. **e**, scDesign3’s model selection functionality allows the evaluation of pseudotime goodness-of-fit using the Bayesian information criterion (BIC). Three pseudotime inference methods—TSCAN, Monocle3, and Slingshot—have BICs evaluated on a synthetic scRNA-seq dataset generated by scDesign3 (based on EMBYRO in Table S2) with true cell pseudotimes. For interpretability, we plot the relative BIC (rBIC) by subtracting the smallest BIC value in **e** so that the rBIC starts from 0. Top: scDesign3 rBIC (calculated without true cell pseudotimes) vs. *R*^2^ between true and inferred pseudotimes (blue: TSCAN; green: Monocle3; orange: Slingshot; black: perturbed true pseudotime as reference, see Methods 1.3.8). The strong negative correlation (Spearman’s rank correlation coefficient *ρ* = −0.95) indicates that scDesign3 BIC measures pseudotime goodness-of-fit effectively. Bottom: visualization of the inferred pseudotime by TSCAN, Monocle3, and Slingshot; Slingshot’s smallest BIC (best goodness-of-fit) agrees with the visualization. **f**, scDesign3’s model alteration functionality allows users to specify the ground truths of cell-type-specific condition effects. In the dataset CONDITION (Table S2), gene *IFI6* is up-regulated in two cell types (CD16+ monocytes and B cells) from control (green) to stimulation (red). With its parametric model, scDesign3 can simulate data where the gene is up-regulated in both cell types (cond++), unchanged in both cell types (cond−−), or only up-regulated in the first cell type CD16+ monocytes (cond+−). **g**, scDesign3’s model alteration functionality allows it to simulate data with or without batch effects. The real dataset (BATCH in Table S2) contains two batches (10x v2 and v3) (left). scDesign3 can preserve the batch effects in its synthetic data (middle: batch+) or generates synthetic data without batch effects (right: batch−). **h**, scDesign3’s model alteration functionality allows it to synthesize null data that do not have cell clusters. The real dataset (ZHENGMIX4 in Table S2) contains two cell types (left). scDesign3 can resemble the two cell types under the alternative hypothesis (*H*_1_) that two cell types exist (middle). In contrast, under the null hypothesis (*H*_0_) that only one cell type exists, scDesign3 can generate a synthetic null dataset that resembles the real data except for the cell type number (right).

## Results

As an overview, we verify scDesign3’s two functionalities—simulation and interpretation—in a sequential manner. First, we show that the scDesign3 model is reasonable from the perspective that its synthetic data well mimic real data given high-quality cell type labels and cell trajectories. Second, under the assumption that the scDesign3 model is reasonable, we show that scDesign3 enables model-based interpretation of real data, including unsupervised assessment of the goodness-of-fit of inferred cell clusters, trajectories, and spatial locations.

### scDesign3 functionality 1: simulation

We verified scDesign3 as a realistic and versatile simulator in four exemplar settings where existing simulators have gaps: (1) scRNA-seq data of continuous cell trajectories, (2) spatial transcriptomics data, (3) single-cell epigenomics data, and (4) single-cell multi-omics data (Fig. 1). Under each setting, we show that the synthetic data of scDesign3 resemble the test data (i.e., left-out real data unused for training), confirming that the scDesign3 model fits well but does not overfit the training data.

In the first setting about continuous cell trajectories, scDesign3 mimics three scRNA-seq datasets containing single or bifurcating cell trajectories (datasets EMBRYO, MARROW, and PANCREAS in Table S2). Fig. 1b–c and Figs. S1–S3c–d show that scDesign3 generates realistic synthetic cells that resemble left-out real cells, as evidenced by high values (≥ 1.75) of mLISI (mean Local Inverse Simpson’s Index), which indicates the degree of similarity between synthetic and real cells and has a lower bound of 1 and a perfect value of 2 [25]. Moreover, scDesign3 preserves eight gene- and cell-specific characteristics, including gene expression mean and variance, gene detection frequency, cell library size, cell-cell distance, cell detection frequency, cell-cell correlation, and, in particular, gene-gene correlation (Figs. S1–S3a–b). Since no existing simulators can generate cells in continuous trajectories by learning from real data, we benchmarked scDesign3 against ZINB-WaVE, muscat, and SPARSIM—three top-performing simulators for generating discrete cell types in previous benchmark studies [14, 15]—and a deep-learning-based simulator scGAN [26]. The results show that scDesign3 outperforms these four simulators in generating more realistic synthetic cells (by achieving higher mLISI values) and in better preserving the eight gene- and cell-specific characteristics, especially cell-cell distances and gene-gene correlations (Fig. 1b–c and Figs. S1–S3). In addition, scDesign3 can output the pseudotime truths of synthetic cells for benchmarking purposes, a functionality unavailable in existing simulators to our knowledge.

In the second setting about spatial transcriptomics, scDesign3 emulates four spatial transcriptomics datasets generated by the 10x Visium and Slide-seq technologies (datasets VISIUM, SLIDE, OVARIAN, and ACINAR in Table S2). First, Fig. 1d–e and Fig. S4 show that scDesign3 recapitulates the expression patterns of spatially variable genes (by achieving high correlations between the corresponding synthetic and real spatial patterns). Second, Figs. S5–S8a–b show that scDesign3 preserves the eight gene- and cell-specific characteristics. Third, Figs. S5–S8c–d use PCA and UMAP embeddings to confirm that the synthetic data of scDesign3 resemble the test data (mLISI values ≥ 1.87). Fourth, scDesign3 mimics spatial transcriptomics data so that each of three prediction algorithms (gradient boosting machine, random forest, and support vector machine) has highly consistent prediction errors (average Pearson correlation > 0.99) between the models trained on real data and scDesign3 synthetic data separately (Fig. S9); moreover, the scDesign3 model can fit complex spatial patterns in less-structured tissues such as cancer tissues (Fig. S10). Notably, in these examples, scDesign3 generates spatial transcriptomics data from spatial locations without cell type annotations (i.e., scDesign3-spatial; see **Methods 1.1.2)**. Figs. S5–S8 show that these synthetic data of scDesign3 are similarly realistic compared to the synthetic data scDesign3 generates under an ideal scenario where annotated cell types are available (i.e., scDesign3-ideal; see **Methods 1.1.2)**. These results confirm scDesign3’s ability to recapitulate cell heterogeneity without needing cell type annotations. Moreover, by fitting a model for spatial transcriptomics data, scDesign3 can estimate a smooth function for every gene’s expected expression levels at spatial locations, a functionality unachievable by existing scRNA-seq simulators.

In addition, when trained on a pair of scRNA-seq data and multi-cell-resolution spatial transcriptomics data (where each spot contains multiple cells), scDesign3 can generate realistic multicell-resolution spatial transcriptomics data with cell-type proportions specified at each spot (i.e., ground truths) (Fig. 1f; Fig. S11a). Using this functionality to benchmark cell-type deconvolution algorithms for spatial transcriptomics data, we found that CARD [27] and RCTD [28] outperformed SPOTlight [29] in estimating the absolute cell-type proportions, though the three algorithms performed similarly well in estimating each cell type’s relative proportions within a tissue slice (Fig.S11b).

In the third setting about single-cell epigenomics, scDesign3 resembles two single-cell chromatin accessibility datasets profiled by the 10x scATAC-seq and sci-ATAC-seq protocols (datasets ATAC and SCIATAC in Table S2). For both protocols, scDesign3 generates realistic synthetic cells (with each cell represented as a vector of genomic regions’ read counts) despite the higher sparsity of single-cell ATAC-seq data compared to scRNA-seq data (Fig. 1g; Fig. 1h left; Figs. S12–S13). Moreover, coupled with our newly proposed read simulator scReadSim [30], scDesign3 extends the simulation of synthetic cells from the count level to the read level, unblocking its application for benchmarking read-level bioinformatics tools (Fig. 1h right).

In the fourth setting about single-cell multi-omics, scDesign3 mimics a CITE-seq dataset (dataset CITE in Table S2) and simulates a multi-omics dataset from separately measured RNA expression and DNA methylation modalities (dataset SCGEM in Table S2). First, scDesign3 resembles the CITE-seq dataset by simultaneously simulating the expression levels of 1000 highly variable genes and 10 surface proteins. Fig. 1i shows that the RNA and protein expression levels of three exemplary surface proteins are highly consistent between the synthetic data of scDesign3 and the test data. Moreover, scDesign3 recapitulates the correlations between the RNA and protein expression levels of the 10 surface proteins (Fig. S14b). Second, scDesign3 simulates a singlecell multi-omics dataset with joint RNA expression and DNA methylation modalities by learning from (1) two single-omics datasets measuring the two modalities separately (Fig. 1j left) and (2) joint low-dimensional embeddings of the two single-omics datasets. This synthetic multiomics dataset preserves the cell trajectory in the two single-omics datasets (Fig. 1j right). The functionality to generate multi-omics data from single-omics data allows scDesign3 to benchmark the computational methods that integrate modalities from unmatched cells [31].

### scDesign3 functionality 2: interpretation

Providing the first universal probabilistic model for single-cell and spatial omics data, scDesign3 has broad applications beyond generating realistic synthetic data. We summarize the prominent applications of the scDesign3 model in three aspects: model parameters, model selection, and model alteration (Fig. 2a).

First, the scDesign3 model has an interpretable parametric structure consisting of genes’ marginal distributional parameters and pairwise gene correlations. In addition to being interpretable, the scDesign3 model is flexible to incorporate cell covariates (such as cell type, pseudotime, and spatial locations) via the use of generalized additive models (see **Methods 1.1.2)**, making the scDesign3 model fit well to various single-cell and spatial omics data—a property confirmed by scDesign3’s realistic simulation in the aforementioned four settings (Fig. 1). The combined interpretability and flexibility enables scDesign3 to estimate the possibly non-linear relationship between every gene’s mean expression and cell covariates, thus allowing statistical inference of gene expression changes between cell types, along cell trajectories (Fig. 2b), and across spatial locations (Fig. 2c).

Besides inferring every gene’s expression characteristics, scDesign3 also estimates pairwise gene correlations conditional on cell covariates, thus providing insights into the possible gene regulatory relationships within each cell type, at a cell differentiation time, or in a spatial region. Specifically, scDesign3 estimates gene correlations by two statistical techniques, Gaussian copula and vine copula, which have complementary advantages (see **Methods 1.1.3)**: Gaussian copula is fast to fit but only outputs a gene correlation matrix; vine copula is slow to fit but outputs a hierarchical gene correlation network (a “vine” with the top layer indicating the most highly correlated genes, i.e., “hub genes”) and thus more interpretable.

As an example application to a dataset containing four human peripheral blood mononuclear cell (PBMC) types (ZHENGMIX4 in Table S2), Fig 2d shows that Gaussian copula reveals similar gene correlation matrices for similar cell types (regulatory T cells vs. naive cytotoxic T cells) and distinct gene correlation matrices for distinct cell types (CD14+ monocytes vs. naive cytotoxic T cells). Moreover, vine copula discovers canonical cell-type marker genes as hub genes: *LYZ* for CD14+ monocytes and *CD79A* for B cells.

Second, scDesign3 outputs the model likelihood, enabling likelihood-based model selection criteria such as Akaike information criterion (AIC) and Bayesian information criterion (BIC). This model selection functionality allows scDesign3 to evaluate the “goodness-of-fit” of a model to data and to compare competing models with the same types of cell covariates. A noteworthy application of this functionality is to evaluate how well does an inferred latent variable (e.g., cell cluster assignment, cell pseudotime, and cell spatial location) describe data, thus enabling us to evaluate inferred cell clusters, pseudotimes, and spatial locations from the goodness-of-fit perspective in the absence of ground truths or external knowledge. Although scDesign3 AIC and BIC rely on the scDesign3 model and do not represent the ground truth, we demonstrate that scDesign3 AIC and BIC are useful “unsupervised” criteria for assessing how well inferred cell clusters, pseudotimes, and spatial locations agree with data under the scDesign3 model.

For cell clustering, we benchmarked scDesign3 BIC against the “supervised” adjusted Rand index (ARI), which requires true cell cluster labels, and a newly proposed unsupervised criterion, clustering deviation index (CDI) [32], on eight datasets with known cell types in a published benchmark study [33]. The results show that scDesign3 BIC has good agreement with ARI (mean Spearman correlation < −0.7) and has better or similar performance compared to CDI’s performance on six out of the eight datasets (Fig. S15b).

For pseudotime inference, scDesign3 BIC is strongly correlated (mean Spearman correlation < −0.7) with the “supervised” *R*^2^, which measures the consistency between the true and inferred (or perturbed) pseudotimes, on multiple synthetic datasets with true pseudotimes (Fig. 2e top; Fig. S15a). Further, scDesign3 BIC agrees with UMAP visualization: compared to TSCAN and Monocle3, the pseudotime inferred by Slingshot has the best (smallest) BIC and best agrees with the low-dimensional representation of the cell manifold (Fig. 2e bottom).

For spatial location inference, we benchmarked scDesign3 AIC against the mean cosine similarity (a supervised metric that measures the similarity between inferred and true spatial locations) using 2 sets of inferred spatial locations and 10 sets of perturbed spatial locations. We find scDesign3 AIC and the mean cosine similarity negatively correlated (mean Spearman correlation ≤ −0.7) on two spatial transcriptomics datasets MOUSE-CORTEX and MOUSE-VISUAL (Table S2), suggesting that scDesign3 AIC is an effective assessment criterion of spatial locations’ goodness-of-fit (Fig. S15c). Note that scDesign3 AIC outperforms BIC in this case, possibly due to the reason that genes’ spatial patterns are complex and thus need complex models.

Third, scDesign3 has a model alteration functionality enabled by its transparent probabilistic modeling and interpretable parameters: given the scDesign3 model parameters estimated on real data, users can alter the model parameters to reflect a hypothesis (i.e., a hypothetical truth) and generate the corresponding synthetic data that bear real data characteristics. Hence, users can flexibly generate synthetic data with varying ground truths for comprehensive benchmarking of computational methods.

We argue that this functionality is a vital advantage scDesign3 has over deep-learning based simulators [26], which cannot be easily altered to reflect a specific hypothesis. We demonstrate how to use this model alteration functionality in three examples. In the first example, scDesign3 generates synthetic data with different cell-type-specific condition effects (Fig. 2f). In the real data (CONDITION in Table S2), gene *IFI6*’s expression is up-regulated after stimulation in both CD16+ monocytes and B cells (Fig. 2f top-left). With scDesign3’s fitted model, users can alter *IFI6*’s mean parameters to make *IFI6*’s expression up-regulated by stimulation in both cell types (Fig. 2f top-right), unchanged by stimulation in both cell types (Fig. 2f bottom-left), or up-regulated by stimulation in CD16+ monocytes only (Fig. 2f bottom-right). In the second example, scDesign3 generates synthetic datasets with or without batch effects (Fig. 2g). Trained on a real dataset (BATCH in Table S2) containing two batches with batch effects (Fig. 2g left), scDesign3’s model, if without alteration, can generate synthetic data retaining the batch effects (Fig. 2g middle), or it can have the batch parameter altered to generate synthetic data without batch effects (Fig. 2g right). In the third example, scDesign3 generates synthetic data under two hypotheses: the null hypothesis (*H*_0_) that only one cell type exists and the alternative hypothesis (*H*_1_) that two cell types exist (Fig. 2h). Given a real dataset (ZHENGMIX4 in Table S2) containing two cell types (Fig. 2h left), the scDesign3 model can be fitted in two ways: under *H*_1_, the model is fitted using the cell type information (Fig. 2h middle); under *H*_0_, the model is fitted by assuming all cells are of one type (Fig. 2h right). The two fitted models can generate the corresponding synthetic data under *H*_1_ and *H*_0_. Particularly, the synthetic data under *H*_0_ can serve as the negative control for benchmarking computational pipelines that use cell clustering to identify the possible existence of cell types.

In summary, scDesign3 is the first omnibus model-based simulator for single-cell and spatial omics data to accommodate different cell states (discrete cell types and continuous cell trajectories), diverse omics features (e.g., gene expression, chromatin accessibility, protein abundance, and DNA methylation), and complex experimental designs (e.g., batches and conditions). Besides generating realistic synthetic data, scDesign3 offers a comprehensive interpretation of real data, thanks to its use of transparent modeling and interpretable parameters. Specifically, scDesign3 estimates the relationship between every feature (e.g., gene) and cell covariates, along with pairwise feature correlations. Moreover, scDesign3 allows likelihood-based model selection to assess the goodness-of-fit of inferred cell clusters, trajectories, and spatial locations output by computational methods. Of course, this unsupervised model-based assessment cannot replace supervised metrics and cannot be used to compare models with different cell covariate types (e.g., cell clusters vs. cell pseudotime). Uniquely, scDesign3 can generate synthetic data under specific hypotheses (e.g., no differential expression, no batch effects, and no cell types) by altering its model parameters. Although the scDesign3 model should not be treated as the true model, its transparent modeling and interpretable parameters can help users explore, alter, and simulate data. Overall, scDesign3 is a multi-functional suite for benchmarking computational methods and interpreting single-cell and spatial omics data.

## Data Availability

All datasets used in the study are publicly available. Supplementary Table S2 lists the datasets from 11 published studies (sources included). The pre-processed datasets are available at: https://doi.org/10.5281/zenodo.7110762.

## Code Availability

The scDesign3 package is available at https://github.com/SONGDONGYUAN1994/scDesign3. The comprehensive tutorials are available at https://songdongyuan1994.github.io/scDesign3/docs/index.html. In the tutorials, we described the input and output formats, model parameters, and exemplary datasets for each functionality of scDesign3. The source code for reproducing the results are available at: https://doi.org/10.5281/zenodo.7110762.

## Acknowledgements

The authors appreciate the comments and feedback from the members of the Junction of Statistics and Biology at UCLA (http://jsb.ucla.edu).

## 1 Methods

### 1.1 The generative model of scDesign3

#### 1.1.1 Mathematical notations of scDesign3’s training data

The training data of scDesign3 contain three matrices: a cell-by-feature matrix (e.g., features are genes or chromatin regions), a cell-by-state-covariate matrix (e.g., cell-state covariates include the cell type, pseudotime, or spatial coordinate), and an optional cell-by-design-covariate matrix (e.g., design covariates include the batch or condition).

Mathematically, first, we denote by 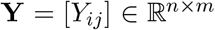 the cell-by-feature matrix with *n* cells as rows, *m* features as columns, and *Y_ij_* as the measurement of feature *j* in cell *i*. For single-cell sequencing data, **Y** is often a count matrix (i.e., 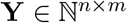, with *Y_ij_* indicating the read or unique molecular identifier (UMI) count of feature *j* in cell *i*); then the sequencing depth (i.e., total number of reads) is 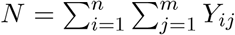.

Second, we denote by 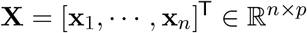 the cell-by-state-covariate matrix with *n* cells as rows and *p* cell-state covariates as columns. Typical cell-state covariates include the cell type (*p* = 1 categorical variable), the cell pseudotime in *p* lineage trajectories (*p* continuous variables), and the 2- or 3-dimensional cell spatial locations (*p* = 2 or 3 continuous variables).

Third, we denote by 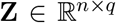 the cell-by-design-covariate matrix with *n* cells as rows and *q* design covariates as columns. Example design covariates are categorical variables such as the batch and condition. Note that **Z** is optional: it is not required if cells are from a single condition and measured in a single batch. To simplify the discussion, in the following text, we write **Z** = [**b**, **c**], where **b** = (*b*_1_,…, *b_n_*)^⊤^ has *b_i_* ∈ {1, · · ·, *B*} representing cell *i*’s batch, and **c** = (*c*_1_,…,*c_n_*)^⊤^ has *c_i_* ∈ {1, · · ·, *C*} representing cell *i*’s condition.

#### 1.1.2 Modeling features’ marginal distributions

For each feature *j* = 1,…, *m* in every cell *i* = 1,…, *n*, the measurement *Y_ij_*—conditional on cell *i*’s state covariates **x_i_** and design covariates **z**_*i*_ = (*b_i_*, *c_i_*)^⊤^—is assumed to follow a distribution *F_j_*(· | **x**_*i*_, **z**_*i*_; *μ_ij_*, *σ_ij_*, *p_ij_*), which is specified as the generalized additive model for location, scale and shape (GAMLSS) [34] (i.e., the distribution family depends on feature *j* only, but the parameters depend on both feature *j* and cell *i*):

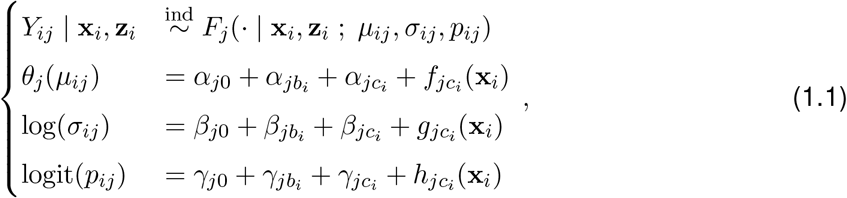

where *θ_j_*(·) denotes feature *j*’s specific link function of the mean parameter *μ_ij_*, depending on *F_j_* (Table S3); *σ_ij_* denotes the scale parameter (e.g., standard deviation or dispersion); *p_ij_* denotes the zero-inflation proportion parameter. Note that *μ_ij_*, *σ_ij_*, and *p_ij_* do not always co-exist, depending on the form of *F_j_* (Table S3). To ensure identifiability, for *j* = 1,…, *m*, we set *α_jb_i__* = *β_jb_i__* = *γ_jb_i__* = 0 when *b_i_* = 1 and *α_jc_i__* = *β_jc_i__* = *γ_jc_i__* = 0 when *c_i_* = 1.

*θ_j_*(*μ_ij_*) is assumed to have feature *j*’s specific intercept *α*_*j*0_, batch *b_i_*’s effect *α_jb_i__* (specific to feature *j*), condition *c_i_*’s effect *α_jc_i__* (specific to feature *j*), and cell-state covariates **x**_*i*_’s effect *f_jc_i__*(**x**_*i*_) (specific to feature *j* and condition *c_i_*).

log(*σ_ij_*) is assumed to have feature *j*’s specific intercept *β*_*j*0_, batch *b_i_*’s effect *β_jb_i__* (specific to feature *j*), condition *c_i_*’s effect *β_jc_i__* (specific to feature *j*), and cell-state covariates **x**_*i*_’s effect *g_jc_i__*(**x**_*i*_) (specific to feature *j* and condition *c_i_*).

logit(*p_ij_*) is assumed to have feature *j*’s specific intercept *γ*_*j*0_, batch *b_i_*’s effect *γ_jb_i__* (specific to feature *j*), condition *c_i_*’s effect *γ_jc_i__* (specific to feature *j*), and cell-state covariates **x**_*i*_’s effect *h_jc_i__*(**x**_*i*_) (specific to feature *j* and condition *c_i_*).

For *θ_j_* (*μ_ij_*), log(*σ_ij_*), and logit(*p_ij_*), the interaction effects are considered between the condition and cell-state covariates, but not between the batch and cell-state covariates. This modeling choice is made based on empirical observations and the simplicity preference [35].

Note that if only the mean parameter *μ_ij_* is assumed to depend on the state covariates **x**_*i*_, batch *b_i_*, and condition *c_i_*, then the GAMLSS degenerates to a generalized additive model (GAM) [36].

Depending on the modality of feature *j* (e.g., a gene’s UMI count), scDesign3 specifies *F_j_* to be one of the six distributions: Gaussian (Normal), Bernoulli, Poisson, Negative Binomial (NB), Zero-inflated Poisson (ZIP), and Zero-inflated Negative Binomial (ZINB); see Table S3 for the specifications. Different specifications of *F_j_* correspond to different link functions *θ_j_*(·) and parameters; see Table S3 for the details.

Depending on cell *i*’s cell-state covariates **x**_*i*_, scDesign3 specifies the functions *f_jc_i__*(·), *g_jc_i__*(·), and *h_jc_i__*(·) in the corresponding forms. See Table S4 for the details. Below are the three typical forms of *f_jc_i__* (·).

1. When the cell-state covariate is the cell type (out of a total of *K_C_* cell types) and **X** = (*x*_1_,…, *x_n_*)^⊤^ with *x_i_* ∈ {1,…, *K_C_*},

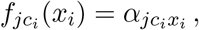

which corresponds to the cell-type *x_i_*’s effect on feature *j* in condition *c_i_*. Note that for identifiability, *αjc_i_x_i_* = 0 if *c_i_* = ^1^.
2. When the cell-state covariates are the cell pseudotimes in *p* lineage trajectories, i.e., **x**_*i*_ = (*x*_*i*1_,…, *x_ip_*)^⊤^ with *x_il_* indicating cell *i*’s pseudotime in the *l*-th lineage trajectory,

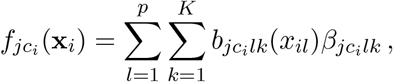

where 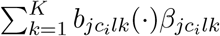 is a cubic spline function for pseudotime in the *l*-th lineage. This formulation means that feature *j* under condition *c_i_* has a specific smooth pattern in lineage *l*. The exact choice *K* is not critical as long as *K* is not too small (see [36]); we set *K* = 10 as default; *K* cannot be larger than the number of data points.
3. When the cell-state covariates are 2-dimensional spatial locations, i.e., **x**_*i*_ = (*x*_*i*1_, *x*_*i*2_)^⊤^ with *x*_*i*1_ and *x*_*i*2_ indicating cell *i*’s spatial locations,

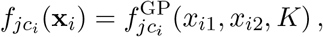

a low-rank Gaussian process smoother described in [36, 37], where *K* is the number of basis functions. This formulation means that feature *j* under condition *c_i_* has a smooth 2-dimensional function (i.e., a surface). The exact choice *K* is not critical as long as *K* is large (see [36]); we set *K* = 400 as default; *K* cannot be larger than the number of data points.

The distribution of (*Y_ij_* | **x**_*i*_, **z**_*i*_) in (1.1) is fitted by the function `gamlss()` in the R package `gamlss` (version 5.4-3) or the function `gam()` in the R package `mgcv` (version 1.8-40). The fitted distribution is denoted as 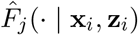, *i* = 1,…, *n*; *j* = 1,…, *m*.

#### 1.1.3 Modeling features’ joint distribution

For cell *i* = 1,…, *n*, we denote its measurements of the *m* features as a random vector **Y**_*i*_ = (*Y*_*i*1_,…, *Y_im_*)^⊤^, whose joint distribution—conditional on cell *i*’s state covariates **x**_*i*_ and design covariates **z**_*i*_—is denoted as 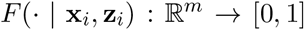. Section 1.1.2 specifies *F_j_*(· | **x**_*i*_, **z**_*i*_), the distribution of (*Y_ij_* | **x**_*i*_, **z**_*i*_), *j* = 1,…, *m*. In scDesign3, the joint cumulative distribution function (CDF) *F*(· | **x**_*i*_, **z**_*i*_) is modeled from the marginal CDFs *F*_1_(· | **x**_*i*_, **z**_*i*_),…, *F_m_*(· | **x**_*i*_, **z**_*i*_) using the copula *C*(· | **x**_*i*_, **z**_*i*_): [0, 1]^m^ → [0, 1]:

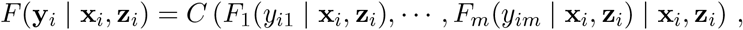

where **y**_*i*_ = (*y*_*i*1_,…, *y_im_*)^⊤^ is a realization of **Y**_*i*_ = (*Y*_*i*1_,…, *Y_im_*)^⊤^.

The copula *C*(· | **x**_*i*_, **z**_*i*_) can be (1) the Gaussian copula or (2) the vine copula, specified below.

The Gaussian copula is defined as

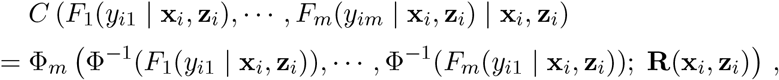

where Φ^−1^ denotes the inverse of the CDF of the standard Gaussian distribution, Φ_*m*_(·; **R**(**x**_*i*_, **z**_*i*_)) denotes the CDF of an *m*-dimensional Gaussian distribution with a zero mean vector and a covariance matrix equal to the correlation matrix **R**(**x**_*i*_, **z**_*i*_).

An issue with the Gaussian copula is that the likelihood calculation is not straightforward in the high-dimensional case when *m* is large and the sample correlation matrix 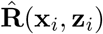, as an estimator of **R**(**x**_*i*_, **z**_*i*_), is not invertible. Then, the likelihood cannot be computed based on 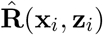. To address this issue, we consider the vine copula.

The vine copula is a way to “decompose” a high-dimensional copula into a sequence of lowdimensional copulas, e.g., bivariate copulas in which every pair of features is modeled as a bivariate Gaussian distribution. In short, the vine copula provides a regular vine (R-vine) structure that uses conditioning to sequentially decompose an *m*-dimensional copula into a sequence of bivariate copulas; then the *m*-dimensional copula density function is approximated by the product of the bivariate copula density functions [38]. The vine copula is advantageous to the Gaussian copula because it enables the likelihood calculation in the high-dimensional case. A detailed definition of the vine copula is in **Supplementary Methods 2**.

To estimate *C*(· | **x**_*i*_, **z**_*i*_) as either the Gaussian or vine copula, we use the plug-in approach that takes the estimated 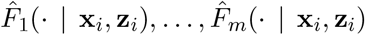 from Section 1.1.2. Specifically, when 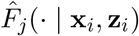 is a continuous distribution, each observed *y_ij_* is transformed as 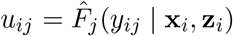. When 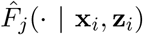 is a discrete distribution with the support on non-negative integers (e.g., the Poisson distribution), since the Gaussian and vine copulas assume that features follow continuous distributions, we use the distributional transformation as in [16]:

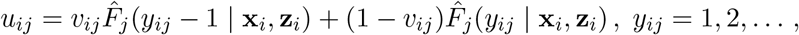

where *v_ij_*’s are sampled independently from Uniform[0, 1], *i* = 1,…, *n*; *j* = 1,…, *m*. To unify and simplify our notations, we write 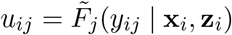, where 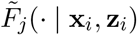 is the CDF of a continuous distribution.

Then *C*(· | **x**_*i*_, **z**_*i*_) is estimated from **u**_1_,…, **u**_*n*_, where **u**_*i*_ = (*u*_*i*1_,…, *u_im_*)^⊤^. For the Gaussian copula, we use the function `cora()` in the R package `Rfast` (version 2.0.6); specifically, 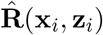 is the sample correlation matrix of {Φ^−1^(**u**_*j*_): (**x**_*j*_, **z**_*j*_) is in a pre-defined neighborhood of (**x**_*i*_, **z**_*i*_)}, where Φ^−1^ (**u**_*i*_) = (Φ^−1^(*u*_*i*1_),…, Φ^−1^ (*u_im_*))^⊤^. For the vine copula, we use the function `vinecop()` in R package `rvinecoplib` (version 0.6.2.1.1).

Then the estimated joint distribution 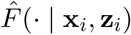 is

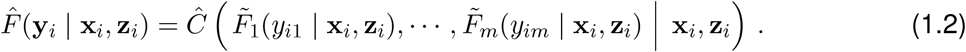

#### 1.1.4 Model likelihood, AIC, and BIC

Given (1.2), the estimated probability density function of cell *i*’s *m*-dimensional feature vector **y**_*i*_, conditional on the cell-state covariates **x**_*i*_ and the design covariates **z**_*i*_, is

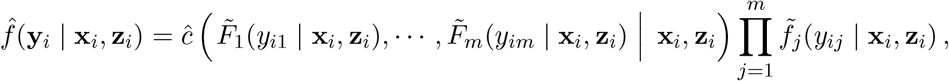

where *ĉ*(· | **x**_*i*_, **z**_*i*_) is the probability density function of *Ĉ*(· | **x**_*i*_, **z**_*i*_), and 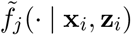 is the probability density function of 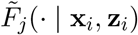. Hence, the log-likelihood is

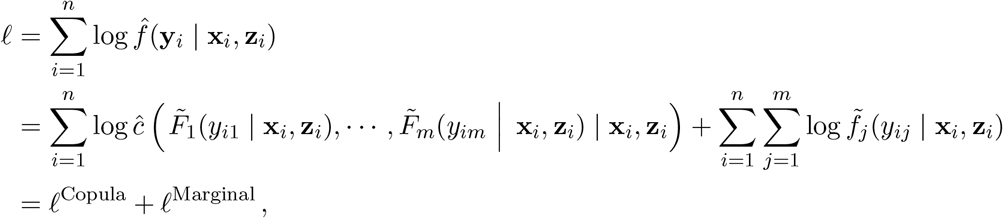

so the log-likelihood can be written as the sum of a copula log-likelihood and a marginal loglikelihood.

Given *k* model parameters and *n* cells (sample size is the number of cells), the Akaike information criterion (AIC) and Bayesian information criterion (BIC) are

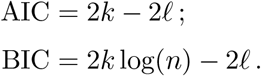

Because of the likelihood decomposition, the AIC and BIC are also decomposable

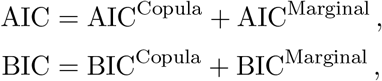

where AIC^Copula^ and BIC^Copula^ only include the number of parameters in *ĉ*(· | **x**_*i*_, **z**_*i*_), and AIC^Marginal^ and BIC^Marginal^ only include the total number of parameters in 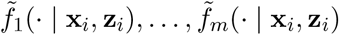.

### 1.2 Synthetic data generation by scDesign3

To generate a synthetic cell-by-feature matrix 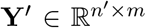, which contains *n′* synthetic cells and the same *m* features as in the training data, scDesign3 allows the specification of a cell-by-state-covariate matrix 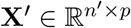 and an optional cell-by-design-covariate matrix 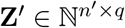 (depending on whether the training data have **Z**) for the *n′* synthetic cells. Note that **X**′ and **Z**′ can be specified by users, generated by resampling the rows of **X** and **Z**, or sampled from some generative models of the rows of **X** and **Z**.

Given **X**, **Z**, and the fitted distributions in Sections 1.1.2 and 1.1.3, scDesign3 samples *n*′ synthetic cells in the following steps.

First, for each synthetic cell *i*′, given its cell-state covariates **x**_*i′*_ and design covariates **z**_*i′*_, we sample its *m*-dimensional probability vector from the *m*-dimensional copula estimated in Section 1.1.3:

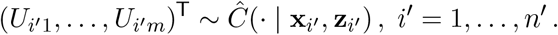

Second, based on the *m* features’ fitted marginal distributions in Section 1.1.2, we calculate the conditional distribution of *Y_i′j_*, the measurement of feature *j* in synthetic cell *i′*, given the synthetic cell’s cell-state covariates **x**_*i′*_ and design covariates **z**_*i′*_ = (*b_i′_*, *c_i′_*)^⊤^, where *b_i′_* ∈ {1,…, *B*} and *c_i′_* ∈ {1,…, *C*}:

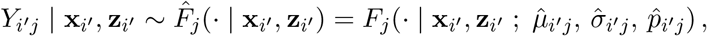

where

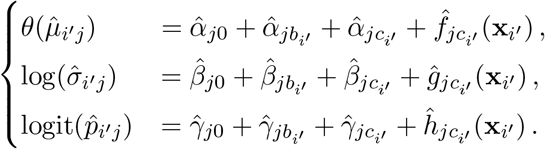

Note that 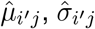, and 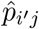 may not be all required, depending on the form of *F_j_* (Table S3).

Then the *m*-dimensional feature vector of synthetic cell *i′* is (*Y*_*i′*1_,…, *Y_i′m_*)^⊤^, where

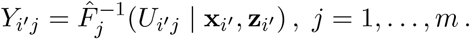

Thanks to the parametric form of 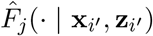, users can generate the synthetic data in their demand by modifying the parameters. For instance, if users want the expected sequencing depth of **Y**′ to change from *N* (the sequencing depth of **Y**) to *N′*, they can scale the mean parameter:

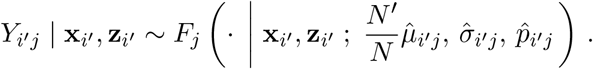

If users want to remove the batch effects, they can set 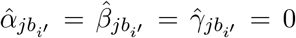, ∀*i′*, *j*. If users want to remove the condition effects, they can set 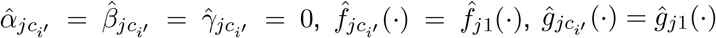, and *ĥ_jc_i′__* (·) = *ĥ*_*j*1_ (·), ∀*i′*, *j*.

### 1.3 Data analysis

#### 1.3.1 Data preprocessing

**Supplementary Table** S2 lists the real datasets from 11 published studies. Since scDesign3 can directly model count data, we did not perform data transformation (e.g., log-transformation) on the cell-by-feature count matrices.

For each cell-by-feature count matrix **Y**, feature screening was used to retain informative features only and save computation time. For every scRNA-seq dataset, we used the R package `scran` (version 1.20.1) [39] to select the top 1000 highly variable genes (HVGs). For the 10x scATAC-seq dataset (ATAC), we used the R package `Signac` (version 1.7.0) [40] to first obtain a cell-by-peak matrix and then select 1133 differentially accessible peaks. For the sci-ATAC-seq data, the preprocessing and feature selection steps are described in [30]. For the 10x Visium datasets (VISIUM, OVARIAN, and ACINAR), we used the R package `Seurat` (version 4.1.1) to select the top 1000 spatially variable genes (SVGs). For the Slide-seq dataset (SLIDE), we selected the top 1000 genes with the smallest *p*-values output by `SPARK-X` [41].

For each dataset, the cell-by-state-covariate matrix **X** was from the original study (if the cellstate covariates are cell types or spatial locations) or inferred by the R package `Slingshot` (version 2.2.1) [42] (if the cell-state covariates are pseudotime values in trajectory lineages).

For each dataset, the optional cell-by-design-covariate matrix **Z** was from the original study if available.

#### 1.3.2 Dimensionality reduction and visualization

To compare scDesign3’s synthetic data with real test data, we used the R package `irlba` (version 2.3.5) to calculate the top 50 principal components (PCs) of the test cell-by-feature matrix (after log-transformation); next, we used the R package `umap` (version 0.2.8.0) to project the test cells from the 50-dimensional PC space to the 2-dimensional UMAP space. Then, we applied the same PCA-UMAP projection to scDesign3’s synthetic cells using the R function `predict()`. Using the same projection ensures that the test cells and synthetic cells are embedded in the same 2-dimensional space and thus comparable.

Unless otherwise noted, the figures were made by the R package `ggplot2` (version 3.3.6). The coverage plot in Fig. 1g was generated by `IGV` (version 2.12.3).

#### 1.3.3 Evaluation metrics

- **mLISI**: To measure the similarity between test cells and synthetic cells in the 2-dimensional space, we used the mean of local inverse Simpson’s index (mLISI) [25] across all cells as the metric. Specifically, if a cell’s neighboring cells are from one group (e.g., test cells or synthetic cells), the cell’s local inverse Simpson’s index (LISI) is 1; otherwise, if a cell’s neighboring cells comprise two groups equally, the cell’s LISI is 2. The mLISI is the average of all cells’ LISIs. Hence, a mLISI close to 2 means that the test cells and synthetic cells are perfectly mixed. The mLISI is calculated by the function `evalIntegration()` in the R package `CellMixS` (version 1.8.0).
- **Pearson correlation**: To measure the similarity between real data and the synthetic data when the cell-state covariates are continuous (e.g., pseudotime, spatial locations, and 2-dimensional UMAP embeddings), we also compared supervised learners trained on the real data and the synthetic data respectively. In detail, for every feature (e.g., gene), we trained a flexible learner, the generalized boosted regression model (GBM), separately on the real data and the synthetic data to predict the feature from cell-state covariates; then, we compared the two GBMs by measuring the Pearson correlation *r* between their predicted values from the synthetic data’s cell-state covariates (note that the cell-state covariates can be replaced by a random sample from the covariate space). An *r* close to 1 means that the two GBMs are similar, that is, the feature’s “relationship” with cell-state covariates is similar in the real data and the synthetic data. If all features have *r* close to 1, the synthetic data resembles the real data. The GBMs were trained using the R package `caret` (version 6.0-93).
- **Summary statistics**: In Supplementary Figures S1–S12, we compared the distributions of eight feature-level, cell-level, feature-pair-level and cell-pair level summary statistics between real data and synthetic data. Note that a feature represents a gene in scRNA-seq and spatial transcriptomics data and a peak in scATAC-seq and sci-ATAC-seq data. The eight summary statistics are

1. mean of log expression (feature-level): a feature’s mean of log expression values across all cells;
2. variance of log expression (feature-level): a feature’s variance of log expression values across all cells;
3. feature detection frequency (feature-level): a feature’s proportion of non-zero values across all cells;
4. feature-feature correlation (feature-pair-level): the correlation between two features’ log expression values across all cells;
5. cell library size on the log scale (cell-level): a cell’s log-transformed total read or UMI count (i.e., log per-cell sequencing depth);
6. cell-cell distance (cell-pair level): the Euclidean distance between two cells in the 50-dim PC space;
7. cell detection frequency (cell-level): a cell’s proportion of non-zero values across all features;
8. cell-cell correlation (cell-pair-level): the correlation between two cells’ log expression values across all features.

Feature-feature correlations were calculated for the top 100 highly expressed features in each real dataset and the corresponding synthetic datasets. To measure the similarity between the real and synthetic correlation matrices, we calculate the Pearson correlation *r* across all 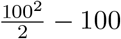 entries in the upper triangle of the correlation matrices.

#### 1.3.4 Boxplots and scatter plots

The boxplots (Fig. 2f) were plotted by the function `geom_boxplot()` in the R package `ggplot2` (version 3.6.6). In each boxplot, the center horizontal line represents the median, the box limits represent the upper and lower quartiles, and the whiskers cover the 1.5× interquartile range. The *p*-value was calculated by the two-sided Wilcoxon rank-sum test.

The scatter plots (Fig. 2e; Fig. S15) were plotted by the function `geom_scatter()` in the R package `ggplot2` (version 3.6.6). In each scatter plot, the *p*-value associated with the Spearman’s correlation coefficient *ρ* was calculated by the one-sided test in the function `stat_cor()` in the R package `ggpubr` (version 0.4.0).

#### 1.3.5 scDesign3’s simulation of spot-resolution transcriptomics data with true cell-type proportions

To generate the synthetic spot-resolution spatial transcriptomics data with true cell-type proportions at each spot, we used a pair of scRNA-seq dataset (MOB-SC) and spatial transcriptomics dataset (MOB-SP) that measured the same biological sample (mouse olfactory bulb). The simulation procedure consists of three steps: the first two steps for parameter estimation from real data and the last step for data simulation.

First, we used scDesign3 to estimate each gene’s mean expression level of each cell type (from scRNA-seq data) and mean expression level at each spatial spot (from spatial transcriptomics data; Fig. S11a Step 1).

Second, using the four cell types’ gene mean expression vectors (one vector per cell type; the cell types are the columns in b; each vector’s elements correspond to genes’ mean expression levels in the cell type) as the reference data and the spatial spots’ gene mean expression vectors (one vector per spot) as the query data, we used the cell-type decomposition method CIBER-SORT [43, 44] to estimate each spot’s cell-type proportions (Fig. 1f left; Fig. S11a Step 2), which we then considered as the spot’s true cell-type proportions in scDesign3’s simulation; note that CIBERSORT could be replaced by other decomposition methods.

Third, we used scDesign3 to generate synthetic scRNA-seq data of the four cell types after training scDesign3 on the real scRNA-seq data. Then we simulated spot-resolutoin transcriptomics data as follows. For each real spot, we sampled 100 cells from the four cell types based on the spot’s true cell-type proportions. Specifically, if the true cell-type proportions are *p*_1_,…, *p*_4_, then the numbers of cells sampled from the four cell types would be drawn from a multinomial distribution Multinomial(100, (*p*_1_,…, *p*_4_)). Then we added the sampled cells’ gene expression vectors and divided the summed vector by 10 to form the spot’s gene expression vector (so every spot corresponds to 10 cells on average, consistent with real data) (Fig. S11a Step 3).

Using the synthetic spot-resolution transcriptomics data, we then benchmarked three spatial transcriptomics deconvolution algorithms: CARD [27], RCTD [28], and SPOTlight [29], using the R packages `CARD` (version 1.0), `spacexr` (version 2.1.6), and `SPOTlight` (version 1.0.1), respectively.

#### 1.3.6 scDesign3’ simulation of a multi-omics dataset from single-omics datasets measuring different modalities

To simulate a multi-omics dataset from real single-omics datasets with unmatched in cells, scDesign3 relies on an integration method that projects single-omics data to a joint low-dimensional space. Then scDesign3 considers each cell’s low-dimensional embedding as the cell covariate in its model.

In Fig. 1j, we used a scRNA-seq dataset and a single-cell methylation dataset with unmatched cells. The two datasets’ cells’ joint low-dimensional embeddings were inferred by the integration method Pamona [45], which could be replaced by other integration methods. Then we trained scDesign3 for each modality (RNA or methylation) using the low-dimensional embeddings of the modality’s corresponding real cells. Finally, using the fitted models (one per modality), we generated a synthetic cell with both modalities from each real cell’s low-dimensional embedding.

#### 1.3.7 scDesign3’s assessment of cell clusters’ goodness-of-fit

To show that scDesign3 can assess clustering goodness-of-fit, we used the 8 datasets from the R package `DuoClustering2018` (version 1.10.0), in which each dataset contains cell type labels (“truth”) and various clustering methods’ results with varying numbers of clusters. The adjusted Rand index (ARI), a “supervised” measure calculated between each clustering result and cell type labels, was used as the benchmark standard. scDesign3’s marginal BIC (Section 1.1.4), an “unsupervised” measure that only uses the clustering result but not the cell type labels, was calculated for each clustering result in each dataset. The models used NB distribution. We used scDesign3’s marginal BIC because we observed that it better captures the clustering goodness-of-fit, while scDesign3’s BIC is dominated by the copula BIC, which largely reflects the number of parameters instead of the clustering quality.

In Fig. S15b, we benchmarked scDesign3’s marginal BIC against the ARI and found them to consistently have negative correlations on the 8 datasets, suggesting that scDesign3’s marginal BIC is an effective assessment measure of clustering goodness-of-fit: a lower BIC indicates better clustering.

#### 1.3.8 scDesign3’s assessment of cell pseudotimes’ goodness-of-fit

To show that scDesign3 can assess pseudotime goodness-of-fit, we used 5 synthetic datasets generated by the R package `dyngen` (version 1.0.3) and 3 synthetic datasets generated by scDesign3; each dataset contains cells’ true pseudotime values (“truth”) ranging from 0 to 1. To generate pseudotime with varying quality, we randomly replaced 0%, 10%, 20%, ···, 100% of truth pseudotime values with randomly sampled values from the Uniform[0, 1] distribution. We also added the inferred pseudotime from R package `slingshot` (version 2.4.0), `monocle3` (version 1.0.0) and `TSCAN` (version 1.34.0). The benchmark standard was the “supervised” *R*^2^ between the true pseudotime values and the perturbed pseudotime values. scDesign3’s marginal BIC (Section 1.1.4), an “unsupervised” measure that only uses the perturbed/inferred pseudotime values but not the true pseudotime values, was calculated for each set of perturbed pseudotime values in each dataset. The models used cubic regression splines and NB distribution. We used scDesign3’s marginal BIC because we observed that it better captures the pseudotime goodness-of-fit, while scDesign3’s BIC is dominated by the copula BIC, which largely reflects the number of parameters instead of the pseudotime goodness-of-fit.

In Fig. S15a, we benchmarked scDesign3’s marginal BIC against the *R*^2^ and found them to consistently have negative correlations on the 8 datasets, suggesting that scDesign3’s marginal BIC is an effective assessment measure of pseudotime goodness-of-fit.

#### 1.3.9 scDesign3’s assessment of inferred spatial locations’ goodness-of-fit

To show that scDesign3 can assess the goodness-of-fit of inferred spatial locations, we used two single-cell resolution spatial transcriptomics datasets from Li et al. [46]. The two datasets contain all cells’ measured spatial locations. Then for each spatial transcriptomics dataset, we treated its cells’ gene expression counts as a “pseudo” scRNA-seq dataset, and we inputted this pseudo scRNA-seq data along with the original spatial transcriptomics dataset into Seurat (version 4.1.1), Tangram (version 1.0.0), and novoSpaRc (version 0.4.3)—as an integration task—to infer the spatial locations of the cells in the “pseudo” scRNA-seq dataset. This approach allowed us to evaluate the inferred spatial locations based on the true spatial locations in the original spatial transcriptomics dataset.

The inferred spatial locations by novoSpaRc contained a large proportion of overlapping locations and thus were not used in our assessment. For Seurat and Tangram, we used each method’s inferred spatial locations along with the original gene expression counts to train scDesign3 (with the NB distribution; Table S3) and calculate the likelihood, marginal AIC, and marginal BIC (Section 1.1.4). Note that we only used the top 100 spatially variable genes defined by Moran’s I statistic to train scDesign3 to save computational time. To evaluate the performance of scDesign3’s unsupervised marginal AIC and BIC, we used the mean cosine similarity, a “supervised” measure that averages all cells’ absolute values of the cosine similarity (for each cell, the cosine similarity is calculated between each cell’s true spatial location and inferred spatial location).

Additionally, for each dataset, we randomly shuffled 0%, 10%, 20%, · · ·, 100% of true spatial locations to obtain spatial locations with varying quality. Similar to the Seurat and Tangram’s inferred spatial locations, we used scDesign3’s marginal AIC and BIC to assess the perturbed spatial locations, using the “supervised” mean cosine similarity as the standard.

In Fig. S15c, we find that scDesign3’ marginal AIC and the mean cosine similarity consistently have negative correlations on the two datasets, suggesting that scDesign3’ marginal AIC is an effective assessment measure of spatial locations’ goodness-of-fit: a lower AIC indicates better spatial goodness-of-fit. Note that AIC outperforms BIC in this case, possibly due to the reason that genes’ spatial patterns are complex and thus need complex models.

#### 1.3.10 Implementation of other simulators

We compared scDesign3 with existing scRNA-seq simulators including scGAN, muscat, SPAR-Sim, and ZINB-WaVE.

- For scGAN, we used the docker and the tutorial the authors provided on scGAN’s GitHub (https://github.com/imsb-uke/scGAN; access date: February 7, 2022) to simulate synthetic data.
- For muscat, we first used the R function `prepSim()` to process the training dataset. Then, we ran the R function `simData()` to simulate a synthetic dataset based on the processed training dataset and cell-level information (such as cell types and experimental conditions) of the training dataset. Both functions are from the R package `muscat` (version 1.6.0).
- For SPARSim, we first used the `SPARSim_create_simulation_parameter` function to obtain the parameters for each group of cells in the training dataset, whose cells were grouped by cell types, experimental conditions, or batches. The 3 required input parameters for the `SPARSim_create_simulation_parameter()` function (`intensity, variability`, and `library_size`) were obtained using the `SPARSim_estimate_intensity()`, `SPARSim_estimate_variability()`, and `SPARSim_estimate_library_size()` functions, respectively, for each cell group. Then, we ran the `SPARSim_simulation()` function with the input parameters from the previous step to generate synthetic data. All functions are from the R package `SPARSim` (version 0.9.5).
- For ZINB-WaVE, we used the `zinbFit()` function from the R package `zinbwave` (version 1.15.3), with the count matrix and cell-type labels as inputs.

## 2 Supplementary Methods

### 2.1 The Vine Copula

An *m*-dimensional copula *C* is a multivariate distribution function composed of *m* Uniform[0, 1] marginal distribution functions. A bivariate copula has *m* = 2. The vine copula provides a regular vine (R-vine) structure that uses conditioning and a tree sequence to decompose an *m*-dimensional copula into bivariate copulas: the *m*-dimensional copula density function is the product of the bivariate copula density functions [38].

We use a triplet 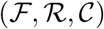 to specify a vine copula, where 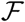 is a vector of marginal distributions, 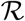 is an R-vine tree sequence, and 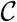 is a set of conditional or unconditional bivariate copulas.

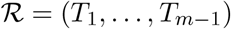 is an R-vine tree sequence if it meets the following conditions [38]:

1. Each tree *T_j_* = (*N_j_*, *E_j_*), where *N_j_* denotes the node set and *E_j_* denotes the edge set, is connected (i.e., there exists an edge path between every two nodes).
2. *T*_1_ is a tree with *m* nodes corresponding to the *m* variables.
3. For all *T_j_* = (*N_j_*, *E_j_*), *j* ≥ 2, *N_j_* = *E*_*j*–1_, i.e., the node set of the *T_j_* is the edge set of *T*_*j*–1_.
4. For all *T_j_*, *j* = 2,…, *m* – 1, any two connected nodes {*n_a_*, *n_b_*} ∈ *E_j_* satisfy |*n_a_* ⋂ *n_b_*| = 1. That is, *n_a_* and *n_b_* are two nodes in *T_j_* and two edges in *T*_*j*–1_; *n_a_* and *n_b_* are connected in *T_j_* if and only if they share a node in *T*_*j*–1_.

After obtaining a valid R-vine tree sequence 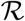, we can construct a unique *m*-dimensional R-vine copula distribution using the triplet 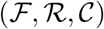 that satisfies the following conditions [47]:

1. Given a random vector **X** = (*X*_1_,…, *X_m_*)^⊤^, 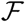 is a vector of the marginal distribution functions of **X**. That is, 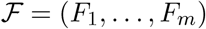, and *F_i_* is continuous and invertible for *i* = 1,…, *m*.
2. 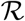 is a R-vine tree sequence specified above.
3. 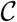 is a set of bivariate copulas, 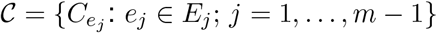, where *C_e_j__* is a bivariate copula corresponding to the edge *e_j_*, and *E_i_* represents the edge set of tree 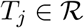.

Here is an example R-vine structure to demonstrate the notations above and the decomposition of an *m*-dimensional joint density function into bivariate copulas. In this example, we have

- **X** = (*X*_1_, *X*_2_, *X*_3_, *X*_4_, *X*_5_)^⊤^ *m* = 5;
- 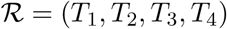;
- *T*_1_ = (*N*_1_, *E*_1_), where *N*_1_ = {*X*_1_, *X*_2_, *X*_3_, *X*_4_, *X*_5_} and *E*_1_ = {(*X*_1_, *X*_2_), (*X*_2_, *X*_3_), (*X*_3_, *X*_4_), (*X*_2_, *X*_5_)};
- *T*_2_ = (*N*_2_, *E*_2_), where *N*_2_ = {(*X*_1_, *X*_2_), (*X*_2_, *X*_3_), (*X*_3_, *X*_4_), (*X*_2_, *X*_5_)} and *E*_2_ = {(*X*_1_, *X*_3_|*X*_2_), (*X*_2_, *X*_4_|*X*_3_), (*X*_3_, *X*_5_|*X*_2_)};
- *T*_3_ = (*N*_3_, *E*_3_), where *N*_3_ = {(*X*_1_, *X*_3_|*X*_2_), (*X*_2_, *X*_4_|*X*_3_), (*X*_3_, *X*_5_|*X*_2_)} and *E*_3_ = {(*X*_1_, *X*_4_|*X*_2_, *X*_3_), (*X*_4_, *X*_5_|*X*_2_, *X*_3_)};
- *T*_4_ = (*N*_4_, *E*_4_), where *N*_4_ = {(*X*_1_, *X*_4_|*X*_2_, *X*_3_), (*X*_4_, *X*_5_|*X*_2_, *X*_3_)} and *E*_4_ = {(*X*_1_, *X*_5_|*X*_2_, *X*_3_, *X*_4_)}.

Then, the joint density of **X** can be written as

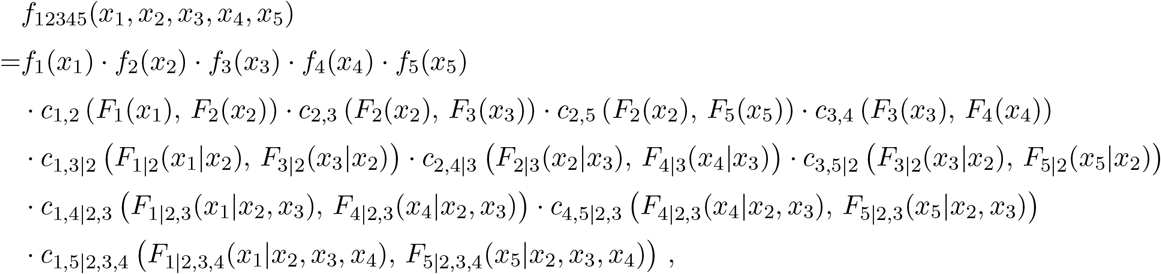

where *c_i,j|D_*: [0, 1]^2^ → [0, ∞) is a bivariate copula density function of *F_i|D_*(*X_i_*) and *F_j|D_*(*X_j_*) conditional on the variable set {*X_k_*: *k* ∈ *D*}, and *F_i|D_* is the conditional CDF of *X_i_* given {*X_k_*: *k* ∈ *D*}, *i* = 1,…, *m*.

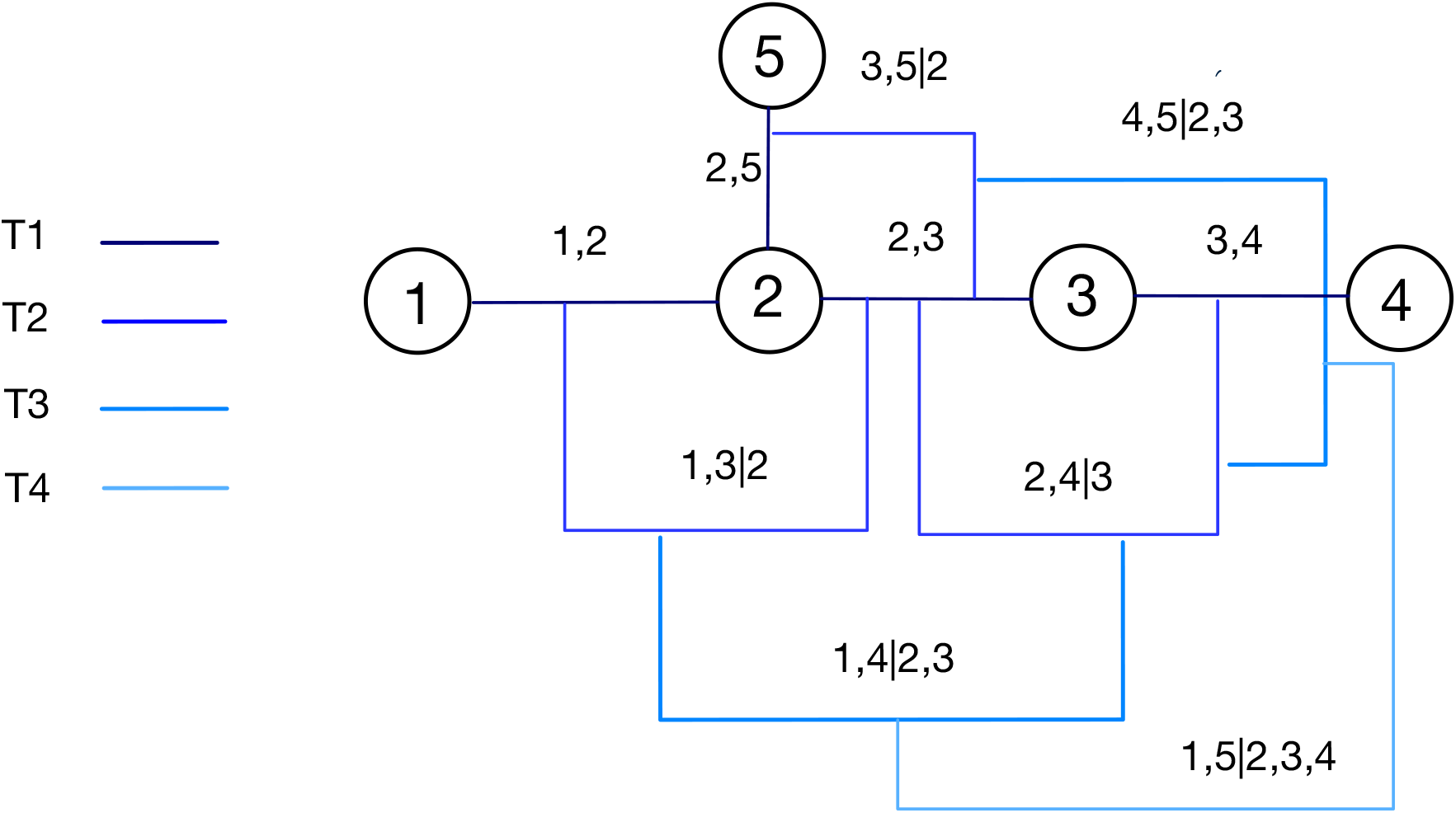

## Supplementary Figures

**Figure S1:**
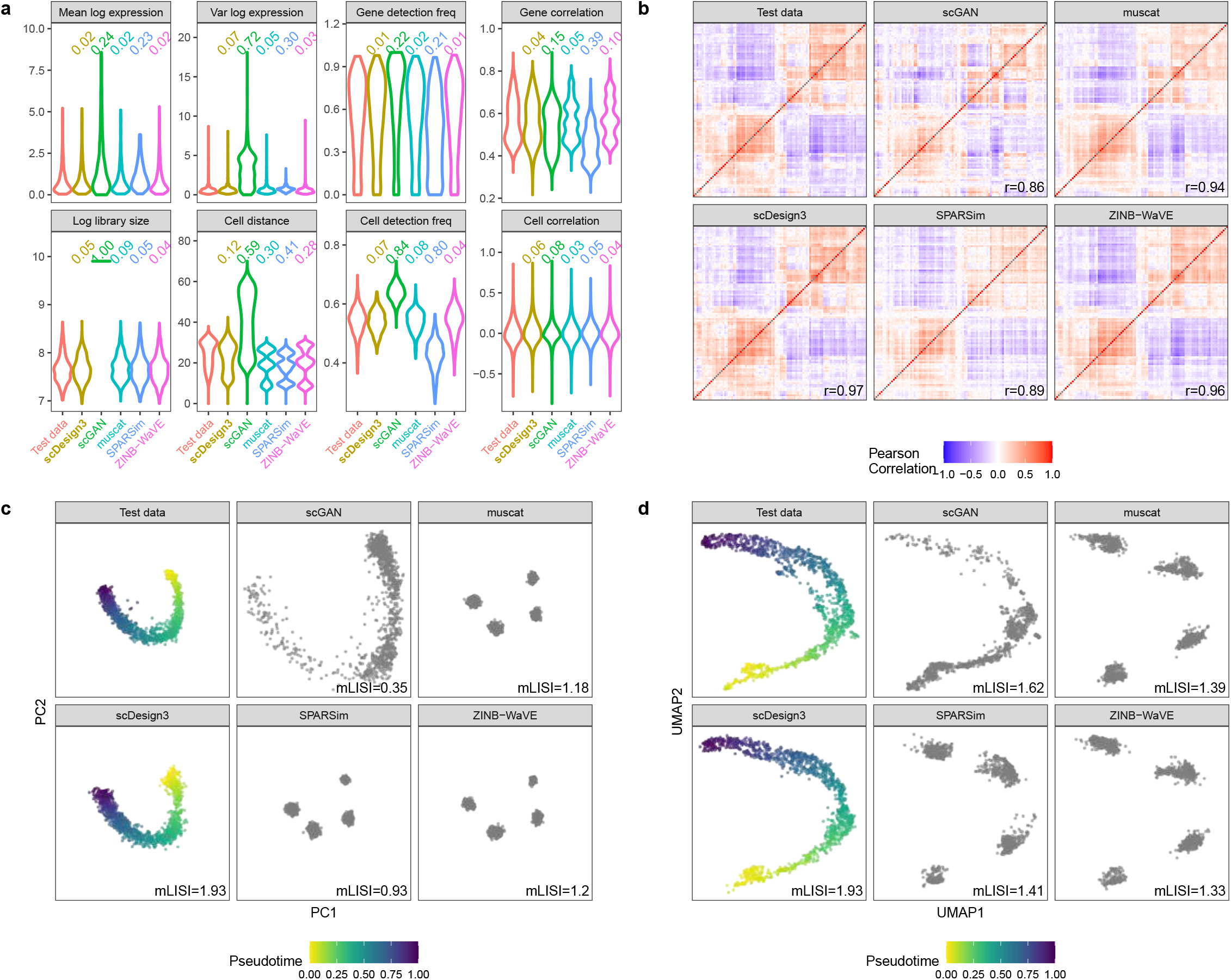
Benchmarking scDesign3 against four existing scRNA-seq simulators (scGAN, muscat, SPARSim, and ZINB-WaVE) for generating scRNA-seq data from a single trajectory (mouse pancreatic endocrinogenesis). **a**, Distributions of eight summary statistics in the test data and the synthetic data generated by scDesign3 and the four simulators. Each number on top of a violin plot (the distribution of a summary statistic in a synthetic dataset) is the Kolmogorov–Smirnov (KS) distance between the synthetic data distribution (indicated by that violin plot) and the test data distribution. A smaller number indicates better agreement between the synthetic data and the test data in terms of that summary statistic’s distribution. **b**, Heatmaps of the gene-gene correlation matrices (showing top 100 highly expressed genes) in the test data and the synthetic data generated by scDesign3 and the four simulators. The Pearson’s correlation coefficient *r* measures the similarity between two correlation matrices, one from the test data and the other from the synthetic data. **c**, PCA visualization (top two PCs) of the test data and the synthetic data generated by scDesign3 and the four simulators. The color labels each cell’s pseudotime value; note that only the synthetic data by scDesign3 outputs the pseudotime truths. An mLISI value close to 2 means that the synthetic data resemble the real data well in the low-dimensional space. **d**, UMAP visualization of the real data and the synthetic data generated by scDesign3 and the four simulators.

**Figure S2:**
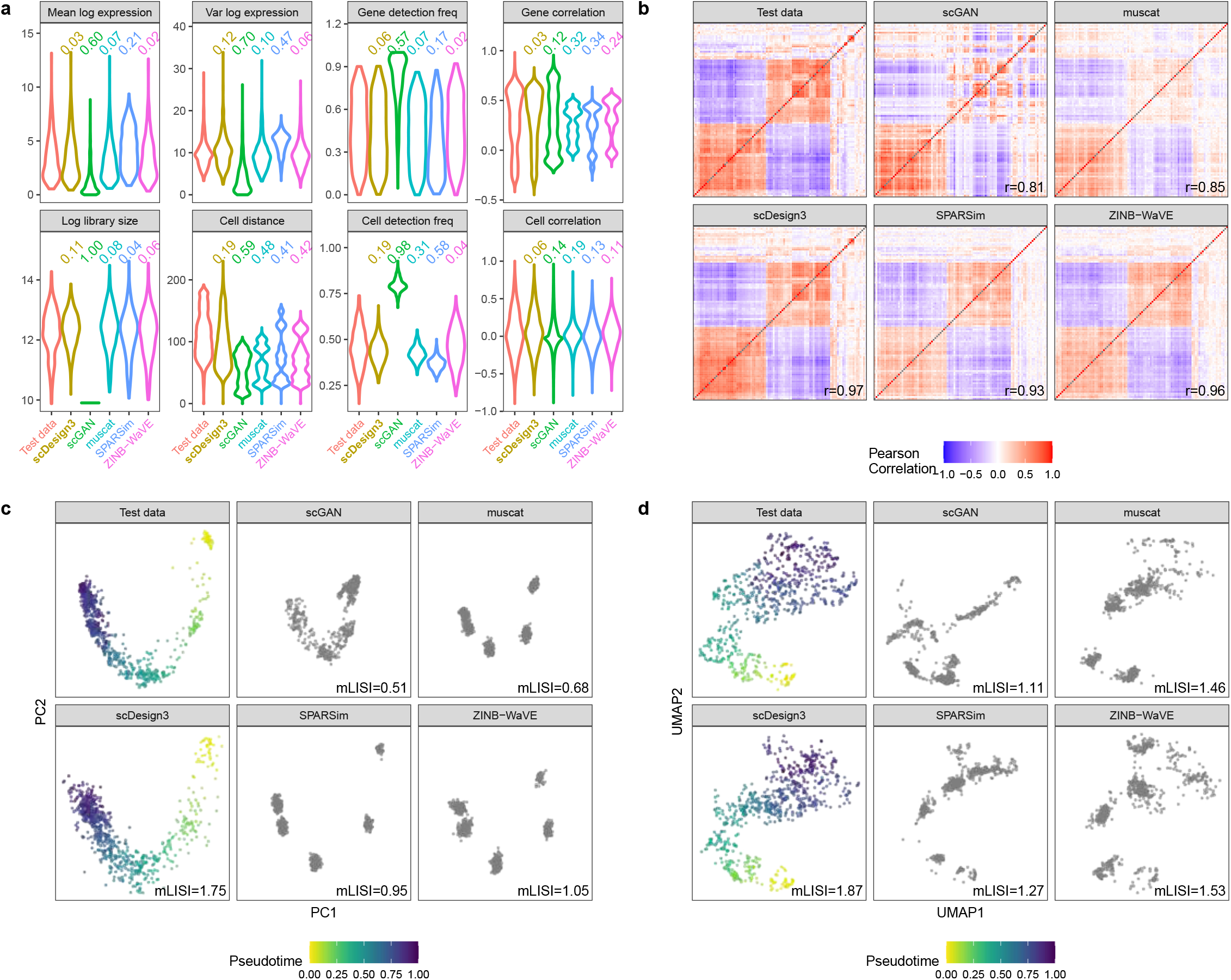
Benchmarking scDesign3 against four existing scRNA-seq simulators (scGAN, muscat, SPARSim, and ZINB-WaVE) for generating scRNA-seq data from a single trajectory (human preimplantation embryos). **a**, Distributions of eight summary statistics in the test data and the synthetic data generated by scDesign3 and the four simulators. Each number on top of a violin plot (the distribution of a summary statistic in a synthetic dataset) is the Kolmogorov–Smirnov (KS) distance between the synthetic data distribution (indicated by that violin plot) and the test data distribution. A smaller number indicates better agreement between the synthetic data and the test data in terms of that summary statistic’s distribution. **b**, Heatmaps of the gene-gene correlation matrices (showing top 100 highly expressed genes) in the test data and the synthetic data generated by scDesign3 and the four simulators. The Pearson’s correlation coefficient *r* measures the similarity between two correlation matrices, one from the test data and the other from the synthetic data. **c**, PCA visualization (top two PCs) of the test data and the synthetic data generated by scDesign3 and the four simulators. The color labels each cell’s pseudotime value; note that only the synthetic data by scDesign3 outputs the pseudotime truths. An mLISI value close to 2 means that the synthetic data resemble the real data well in the low-dimensional space. **d**, UMAP visualization of the real data and the synthetic data generated by scDesign3 and the four simulators.

**Figure S3:**
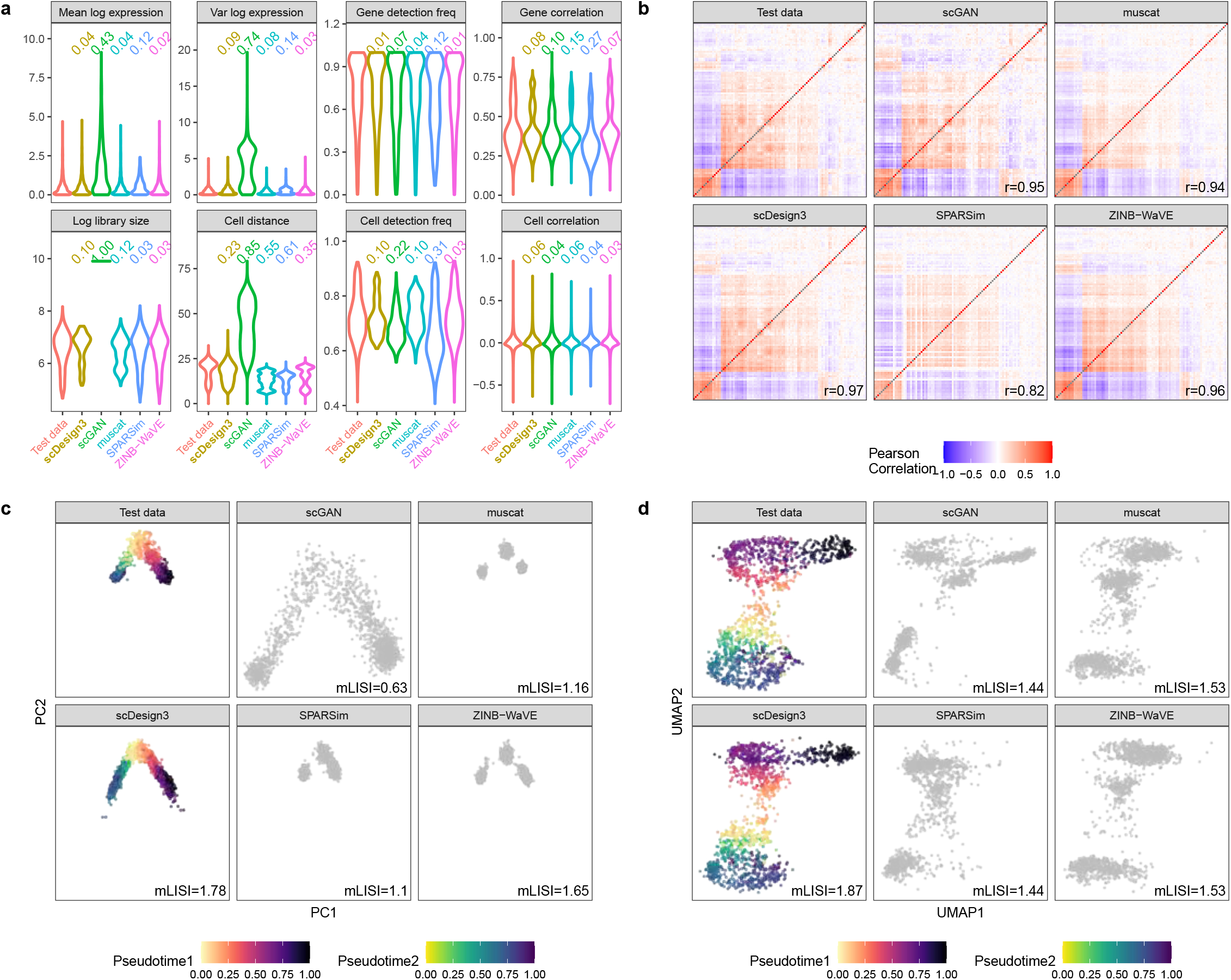
Benchmarking scDesign3 against four existing scRNA-seq simulators (scGAN, muscat, SPARSim, and ZINB-WaVE) for generating scRNA-seq data from bifurcating trajectories (myeloid progenitors in mouse bone marrow). **a**, Distributions of eight summary statistics in the test data and the synthetic data generated by scDesign3 and the four simulators. Each number on top of a violin plot (the distribution of a summary statistic in a synthetic dataset) is the Kolmogorov–Smirnov (KS) distance between the synthetic data distribution (indicated by that violin plot) and the test data distribution. A smaller number indicates better agreement between the synthetic data and the test data in terms of that summary statistic’s distribution. **b**, Heatmaps of the gene-gene correlation matrices (showing top 100 highly expressed genes) in the test data and the synthetic data generated by scDesign3 and the four simulators. The Pearson’s correlation coefficient *r* measures the similarity between two correlation matrices, one from the test data and the other from the synthetic data. **c**, PCA visualization (top two PCs) of the test data and the synthetic data generated by scDesign3 and the four simulators. The color labels each cell’s pseudotime value; note that only the synthetic data by scDesign3 outputs the pseudotime truths. An mLISI value close to 2 means that the synthetic data resemble the real data well in the low-dimensional space. **d**, UMAP visualization of the real data and the synthetic data generated by scDesign3 and the four simulators.

**Figure S4:**
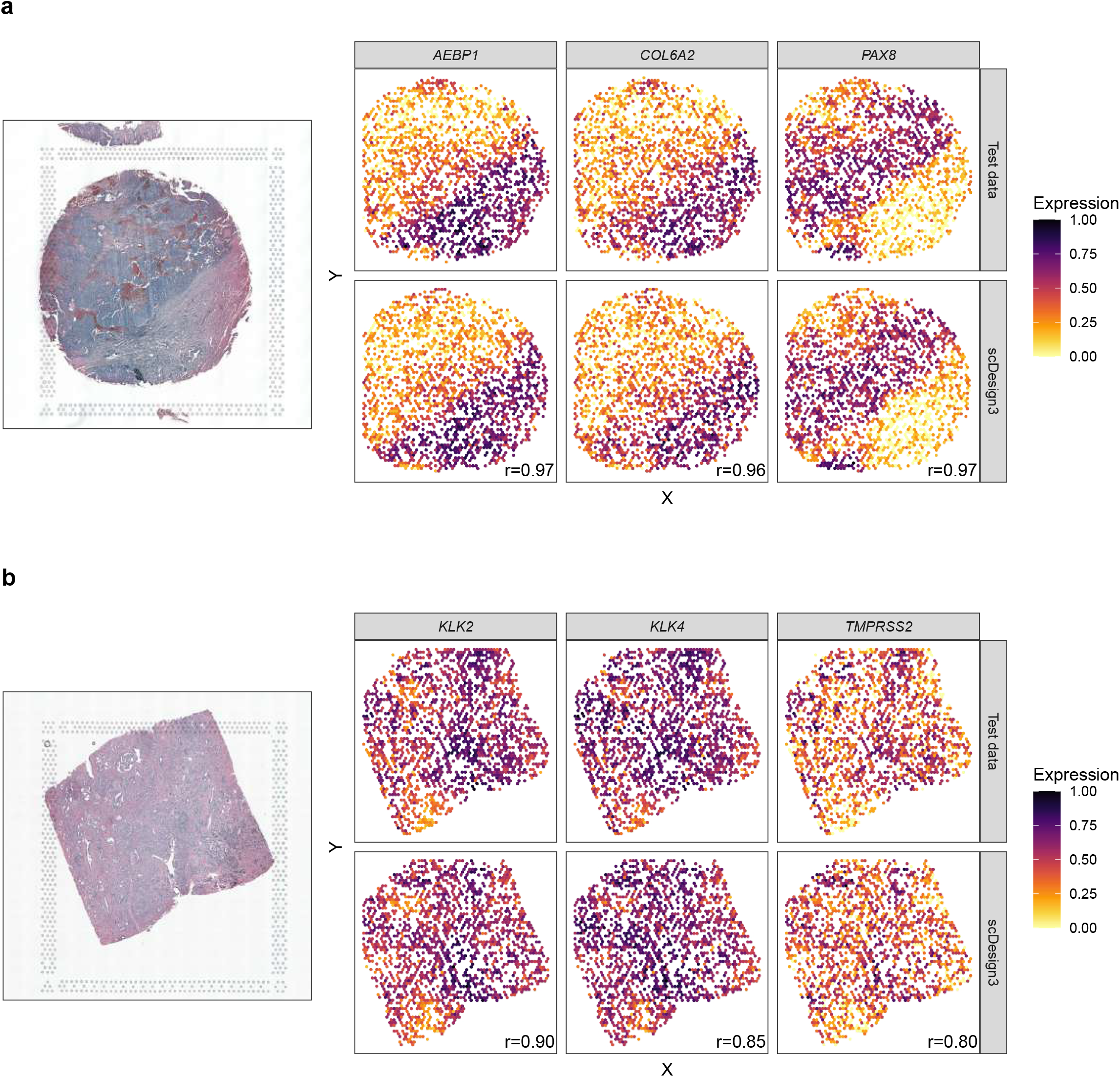
scDesign3 simulates realistic gene expression patterns for cancer transcriptomics datasets: human ovarian cancer (a) and human prostate cancer, acinar cell carcinoma (b). The tissue samples are measured with both H&E (hematoxylin and eosin stain, left) and spatial transcriptomics (right, three cancer-related genes). Large Pearson correlation coefficients (r) represent similar spatial patterns in synthetic and test data.

**Figure S5:**
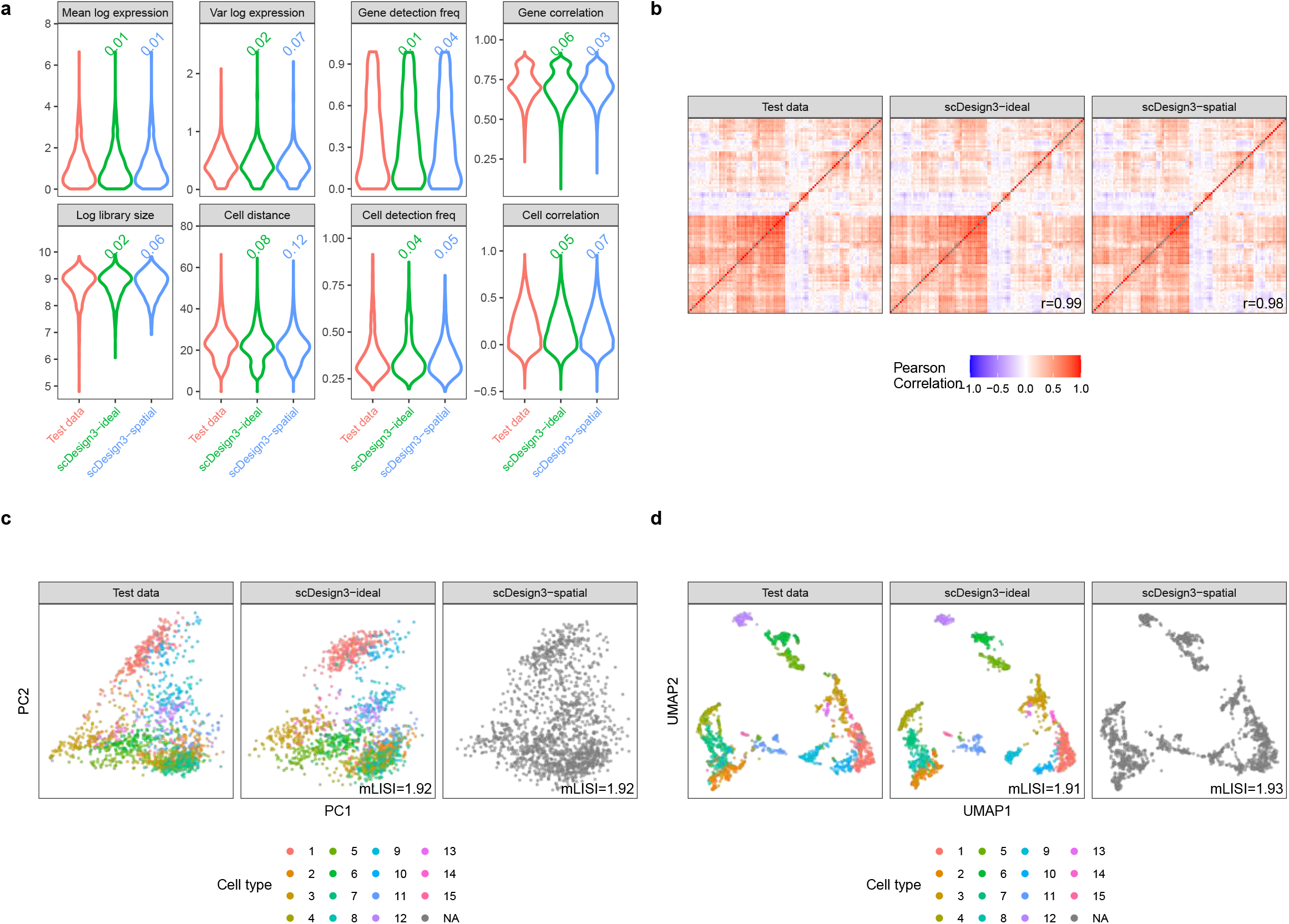
scDesign3 simulates 10x Visium spatial transcriptomics data (sagital mouse brain slices). **a**, Distributions of eight summary statistics in the test data and the synthetic data generated by scDesign3 using cell type labels (scDesign3-ideal) and spatial locations (scDesign3-spatial), respectively. Each number on top of a violin plot (the distribution of a summary statistic in a synthetic dataset) is the Kolmogorov–Smirnov (KS) distance between the synthetic data distribution (indicated by that violin plot) and the test data distribution. A smaller number indicates better agreement between the synthetic data and the test data in terms of that summary statistic’s distribution. **b**, Heatmaps of the gene-gene correlation matrices (showing top 100 highly expressed genes) in the test data and the synthetic data generated by scDesign3-ideal and scDesign3-spatial. The Pearson’s correlation coefficient *r* measures the similarity between two correlation matrices, one from the test data and the other from the synthetic data. **c**, PCA visualization (top two PCs) of the real data and the synthetic data generated by scDesign3-ideal and scDesign3-spatial. The color labels each cell’s cell type (cluster). Since the scDesgin3-spatial data only uses spatial locations, it does not rely on cell types. An mLISI value close to 2 means that the synthetic data resemble the real data well in the low-dimensional space. **d**, UMAP visualization of the real data and the synthetic data generated by scDesign3-ideal and scDesign3-spatial. In summary, scDesign3 realistically simulates 10x Visium data based on spatial locations without needing cell type annotations.

**Figure S6:**
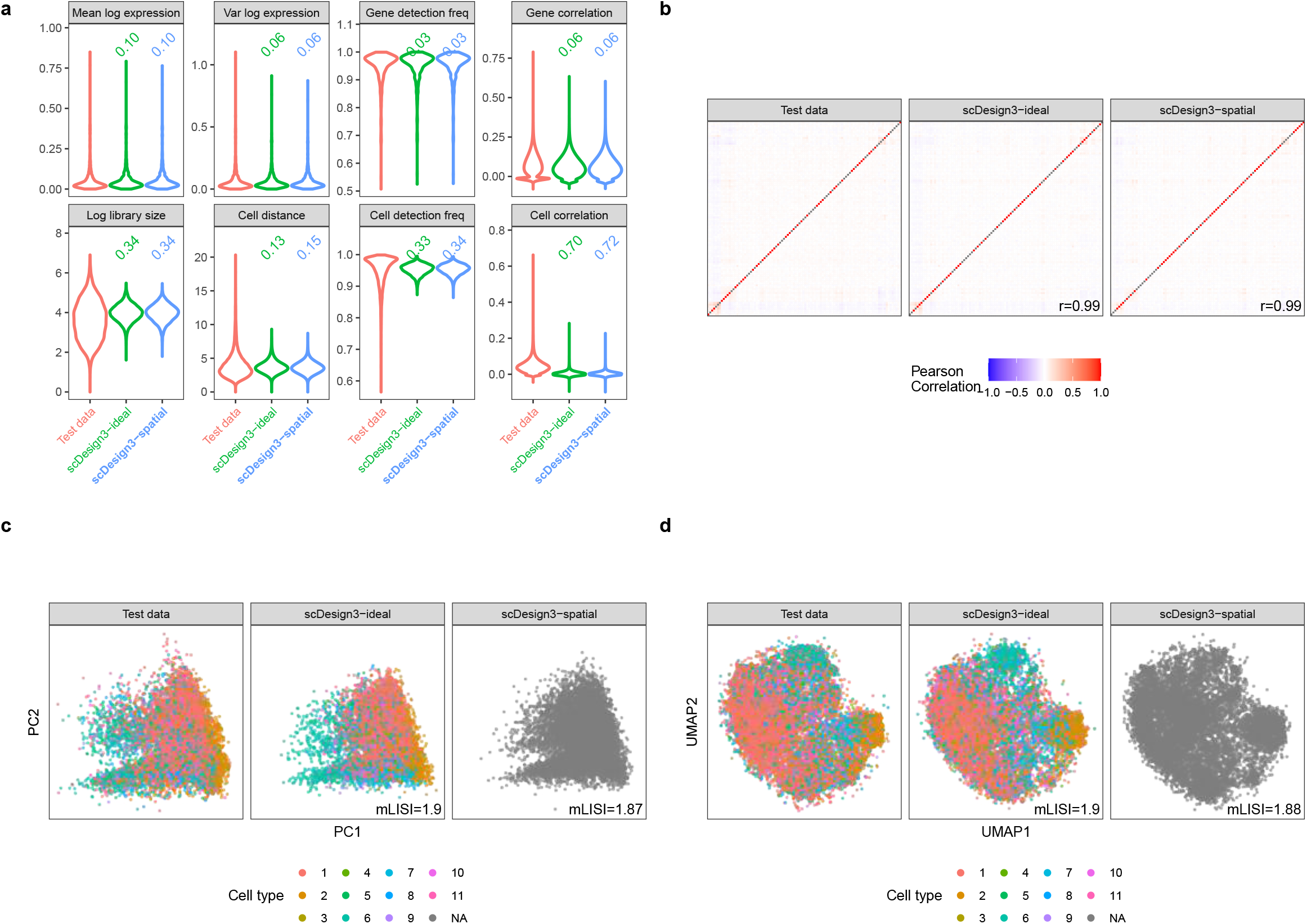
scDesign3 simulates Slide-seq spatial transcriptomics data (coronal cerebellum). **a**, Distributions of eight summary statistics in the test data and the synthetic data generated by scDesign3 using cell type labels (scDesign3-ideal) and spatial locations (scDesign3-spatial), respectively. Each number on top of a violin plot (the distribution of a summary statistic in a synthetic dataset) is the Kolmogorov–Smirnov (KS) distance between the synthetic data distribution (indicated by that violin plot) and the test data distribution. A smaller number indicates better agreement between the synthetic data and the test data in terms of that summary statistic’s distribution. **b**, Heatmaps of the genegene correlation matrices (showing top 100 highly expressed genes) in the test data and the synthetic data generated by scDesign3-ideal and scDesign3-spatial. The Pearson’s correlation coefficient *r* measures the similarity between two correlation matrices, one from the test data and the other from the synthetic data. **c**, PCA visualization (top two PCs) of the real data and the synthetic data generated by scDesign3-ideal and scDesign3-spatial. The color labels each cell’s cell type (cluster). Since scDesgin3-spatial only uses spatial locations, it does not rely on cell types. An mLISI value close to 2 means that the synthetic data resemble the real data well in the low-dimensional space. **d**, UMAP visualization of the real data and the synthetic data generated by scDesign3-ideal and scDesign3-spatial. In summary, scDesign3 realistically simulates Slide-seq data based on spatial locations without needing cell type annotations.

**Figure S7:**
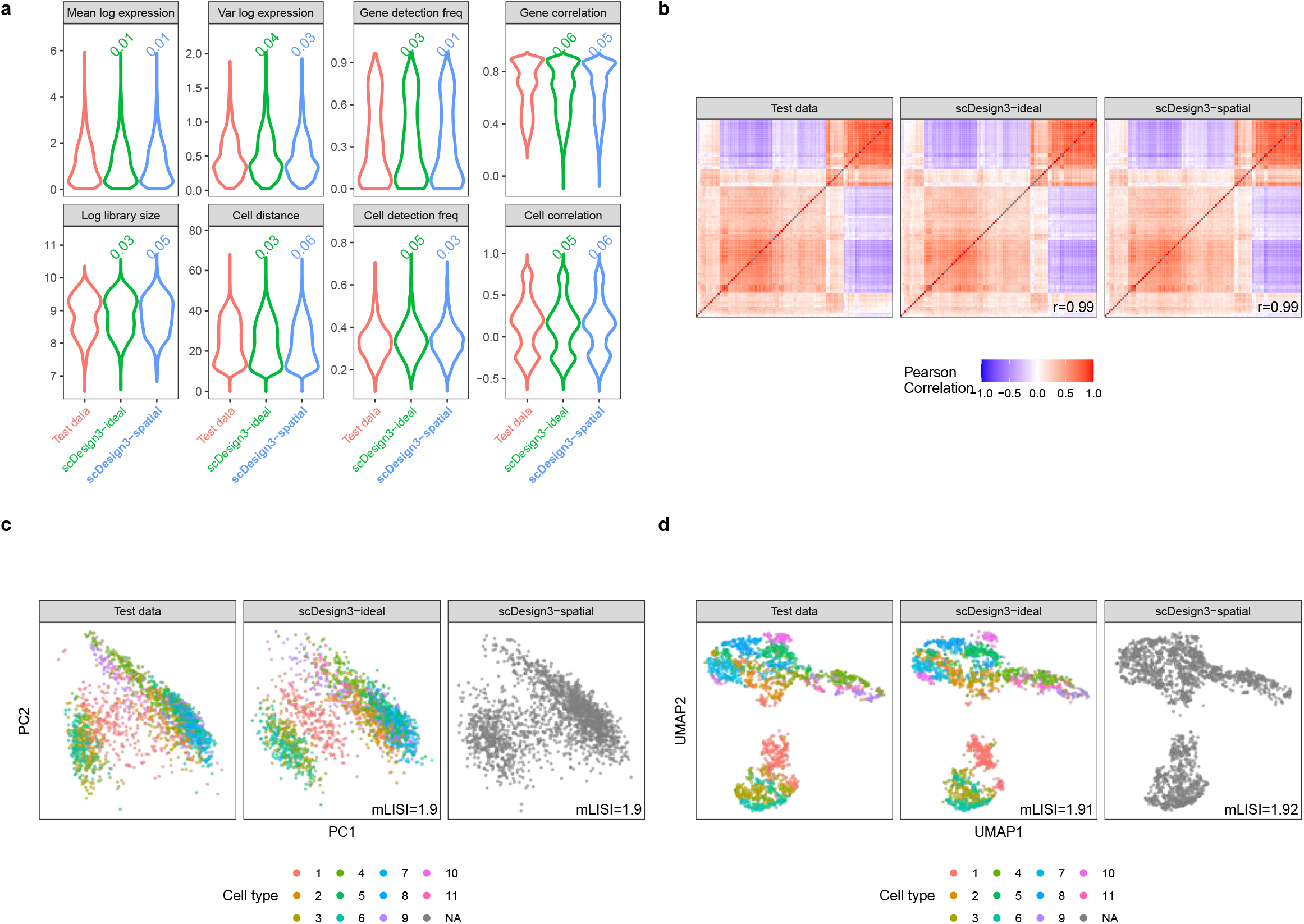
scDesign3 simulates 10x Visium cancer spatial transcriptomics data (human ovarian cancer). **a**, Distributions of eight summary statistics in the test data and the synthetic data generated by scDesign3 using cell type labels (scDesign3-ideal) and spatial locations (scDesign3-spatial), respectively. Each number on top of a violin plot (the distribution of a summary statistic in a synthetic dataset) is the Kolmogorov–Smirnov (KS) distance between the synthetic data distribution (indicated by that violin plot) and the test data distribution. A smaller number indicates better agreement between the synthetic data and the test data in terms of that summary statistic’s distribution. **b**, Heatmaps of the gene-gene correlation matrices (showing top 100 highly expressed genes) in the test data and the synthetic data generated by scDesign3-ideal and scDesign3-spatial. The Pearson’s correlation coefficient *r* measures the similarity between two correlation matrices, one from the test data and the other from the synthetic data. **c**, PCA visualization (top two PCs) of the real data and the synthetic data generated by scDesign3-ideal and scDesign3-spatial. The color labels each cell’s cell type (cluster). Since the scDesgin3-spatial data only uses spatial locations, it does not rely on cell types. An mLISI value close to 2 means that the synthetic data resemble the real data well in the lowdimensional space. **d**, UMAP visualization of the real data and the synthetic data generated by scDesign3-ideal and scDesign3-spatial. In summary, scDesign3 realistically simulates 10x Visium data based on spatial locations without needing cell type annotations.

**Figure S8:**
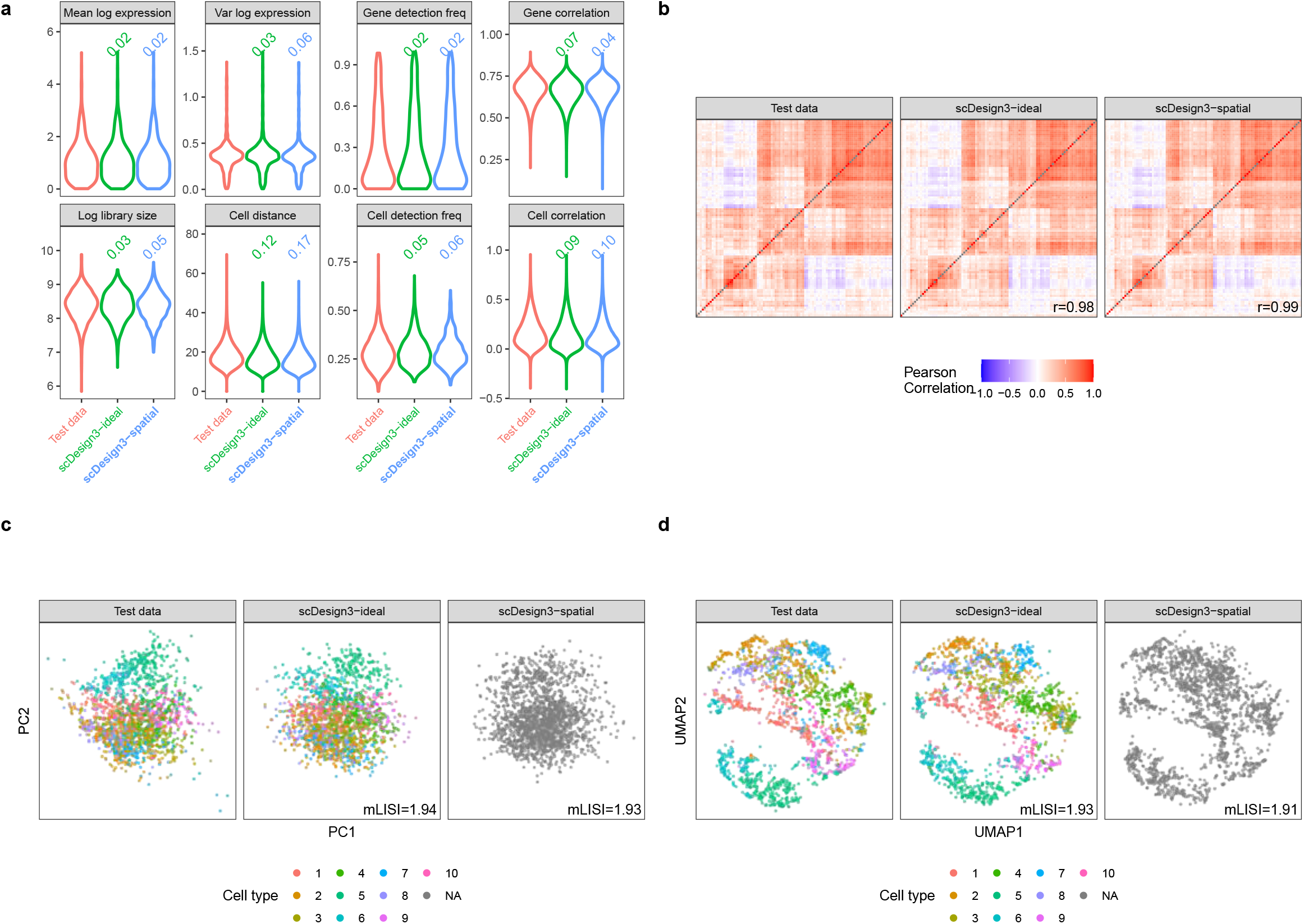
scDesign3 simulates 10x Visium cancer spatial transcriptomics data (human prostate cancer, acinar cell carcinoma). **a**, Distributions of eight summary statistics in the test data and the synthetic data generated by scDesign3 using cell type labels (scDesign3-ideal) and spatial locations (scDesign3-spatial), respectively. Each number on top of a violin plot (the distribution of a summary statistic in a synthetic dataset) is the Kolmogorov–Smirnov (KS) distance between the synthetic data distribution (indicated by that violin plot) and the test data distribution. A smaller number indicates better agreement between the synthetic data and the test data in terms of that summary statistic’s distribution. **b**, Heatmaps of the gene-gene correlation matrices (showing top 100 highly expressed genes) in the test data and the synthetic data generated by scDesign3-ideal and scDesign3-spatial. The Pearson’s correlation coefficient *r* measures the similarity between two correlation matrices, one from the test data and the other from the synthetic data. **c**, PCA visualization (top two PCs) of the real data and the synthetic data generated by scDesign3-ideal and scDesign3-spatial. The color labels each cell’s cell type (cluster). Since the scDesgin3-spatial data only uses spatial locations, it does not rely on cell types. An mLISI value close to 2 means that the synthetic data resemble the real data well in the lowdimensional space. **d**, UMAP visualization of the real data and the synthetic data generated by scDesign3-ideal and scDesign3-spatial. In summary, scDesign3 realistically simulates 10x Visium data based on spatial locations without needing cell type annotations.

**Figure S9:**
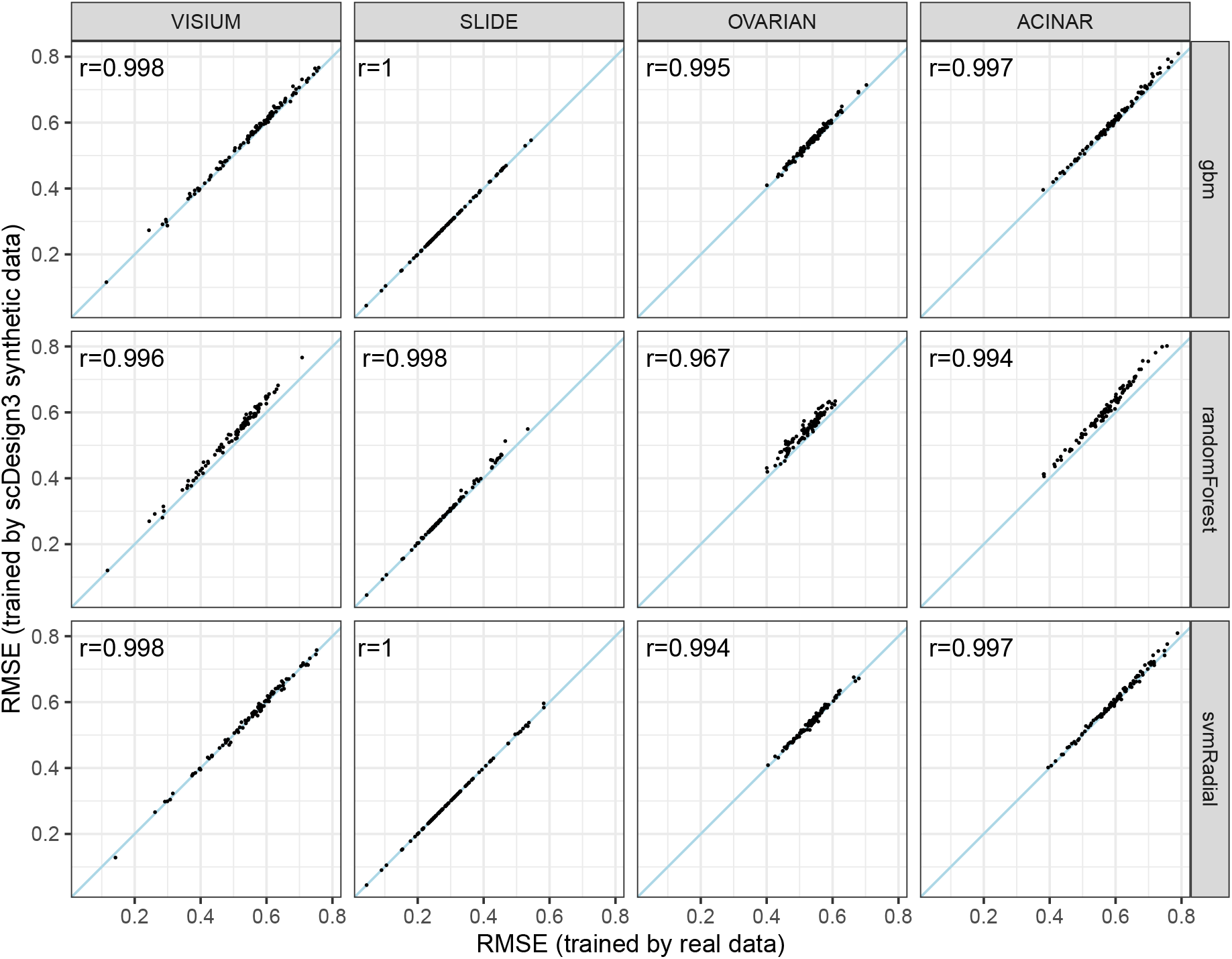
scDesign3 mimics spatial transcriptomics data so that prediction algorithms have similar prediction performance when trained on real data or scDesign3 synthetic data. In detail, we first split each of four spatial transcriptomics datasets (VISIUM, SLIDE, OVARIAN, and ACINAR) into two datasets (training and testing) by randomly splitting the spatial locations into two halves. Second, we use each of the four training datasets to fit scDesign3 and generate the corresponding synthetic dataset. Third, on each pair of training dataset and synthetic dataset (among a total of four pairs), we train each of three prediction algorithms (gbm: gradient boosting machine; randomForest: random forest; svmRadial: support vector machine with the radial kernel) to predict each gene’s expression at a spatial location (input: spatial location; output: the gene’s log(count+1) expression level at the location), obtaining a pair of prediction models for each gene. Fourth, we apply each pair of prediction models to the corresponding testing dataset and calculate each model’s root-mean-squared error (RMSE) for predicting each gene, obtaining a pair of RMSEs. As a result, in each panel, we plot the RMSEs for each prediction algorithm (row) and dataset (column), with each dot in the panel representing a gene. We observe that all genes’ RMSEs are highly similar, reflecting that scDesign3 synthetic data well mimic real data.

**Figure S10:**
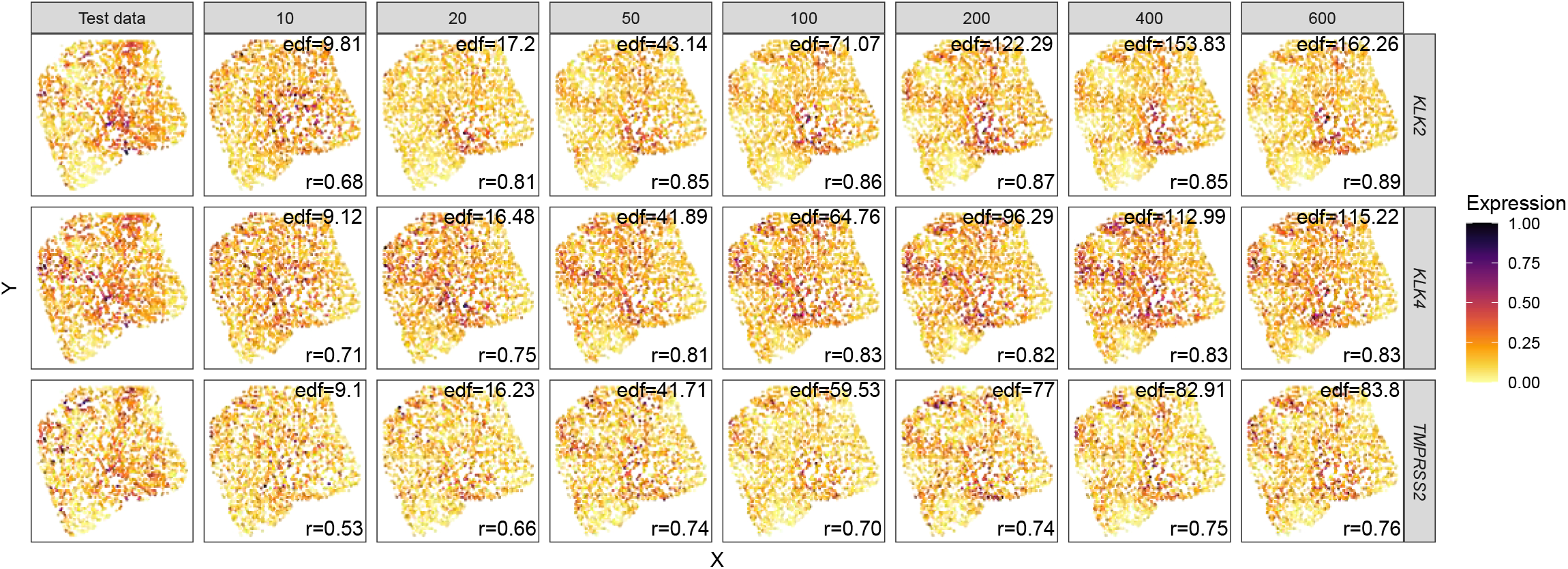
The effect of *K* on simulating spatial transcriptomics data. The rows represent three cancer-related genes; column 1 represents real test data; columns 2–8 represent scDesign3 synthetic data generated using varying input basis numbers *K*. A large Pearson correlation coefficient (*r*) represents similar spatial patterns in synthetic and test data. The effective degrees of freedom (edf) represent the wiggliness of the fitted surface. With a larger *K*, scDesign3 is able to fit more complex patterns. The overfitting issue is accounted for by the automatic smoothness estimation [36]: when *K* is sufficiently large, edf (model complexity) and *r* (model goodness-of-fit) both become stable.

**Figure S11:**
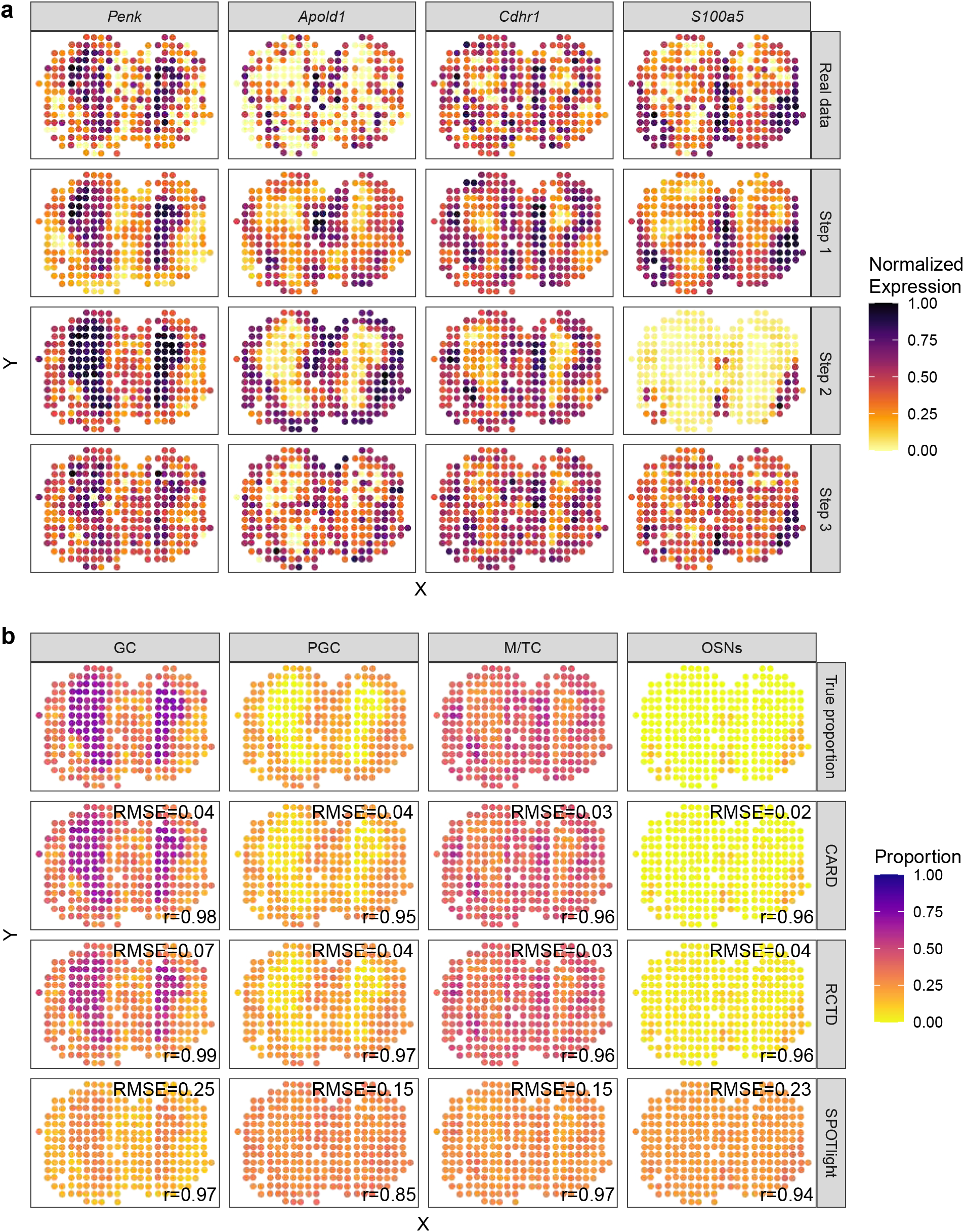
scDesign3 simulates spot-resolution spatial transcriptomics data for benchmarking cell-type deconvolution methods. **a**, the scDesign3 spot simulation mimics the real data well by showing similar expression patterns for the four cell-type marker genes. **b**, Using scDesign3 synthetic data, we benchmark three spatial deconvolution methods (CARD [28], RCTD [27], and SPOTlight [29]). For each of four cell types (columns), we use two metrics—Pearson correlation (*r*) and root-mean-square error (RMSE)—to compare the estimated proportions by each deconvolution method (rows 2–4) to the true proportions (top row). Large *r* values represent similar spatial patterns of proportions, while small RMSE values represent similar values of proportions. Although all three methods well capture the spatial patterns of each cell type’s proportions (evidenced by large *r* values), CARD and RCTD outperform SPOTlight by estimating celltype proportions more accurately (evidenced by smaller RMSE values).

**Figure S12:**
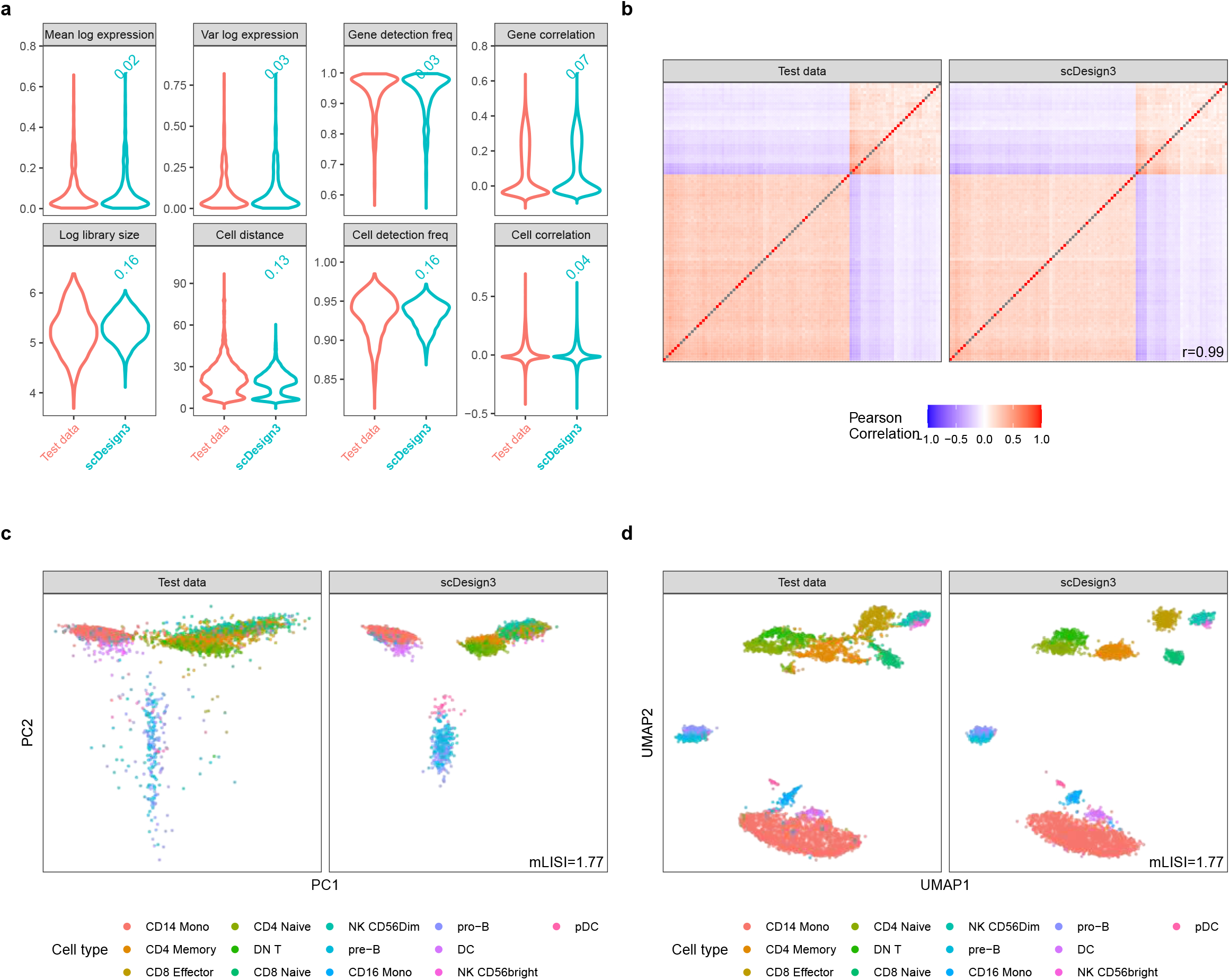
scDesign3 simulates scATAC-seq data (human PBMCs). **a**, Distributions of eight summary statistics in the test data and the synthetic data generated by scDesign3 using cell type labels. Each number on top of a violin plot (the distribution of a summary statistic in a synthetic dataset) is the Kolmogorov–Smirnov (KS) distance between the synthetic data distribution (indicated by that violin plot) and the test data distribution. A smaller number indicates better agreement between the synthetic data and the test data in terms of that summary statistic’s distribution. **b**, Heatmaps of the peak-peak correlation matrices in the test data and the synthetic data generated by scDesign3. The Pearson’s correlation coefficient *r* measures the similarity between two correlation matrices, one from the test data and the other from the synthetic data. **c**, PCA visualization (top two PCs) of the test data and the synthetic data generated by scDesign3. The color labels each cell’s cell type. An mLISI value close to 2 means that the synthetic data resemble the test data well in the low-dimensional space. **d**, UMAP visualization of the test data and the synthetic data generated by scDesign3.

**Figure S13:**
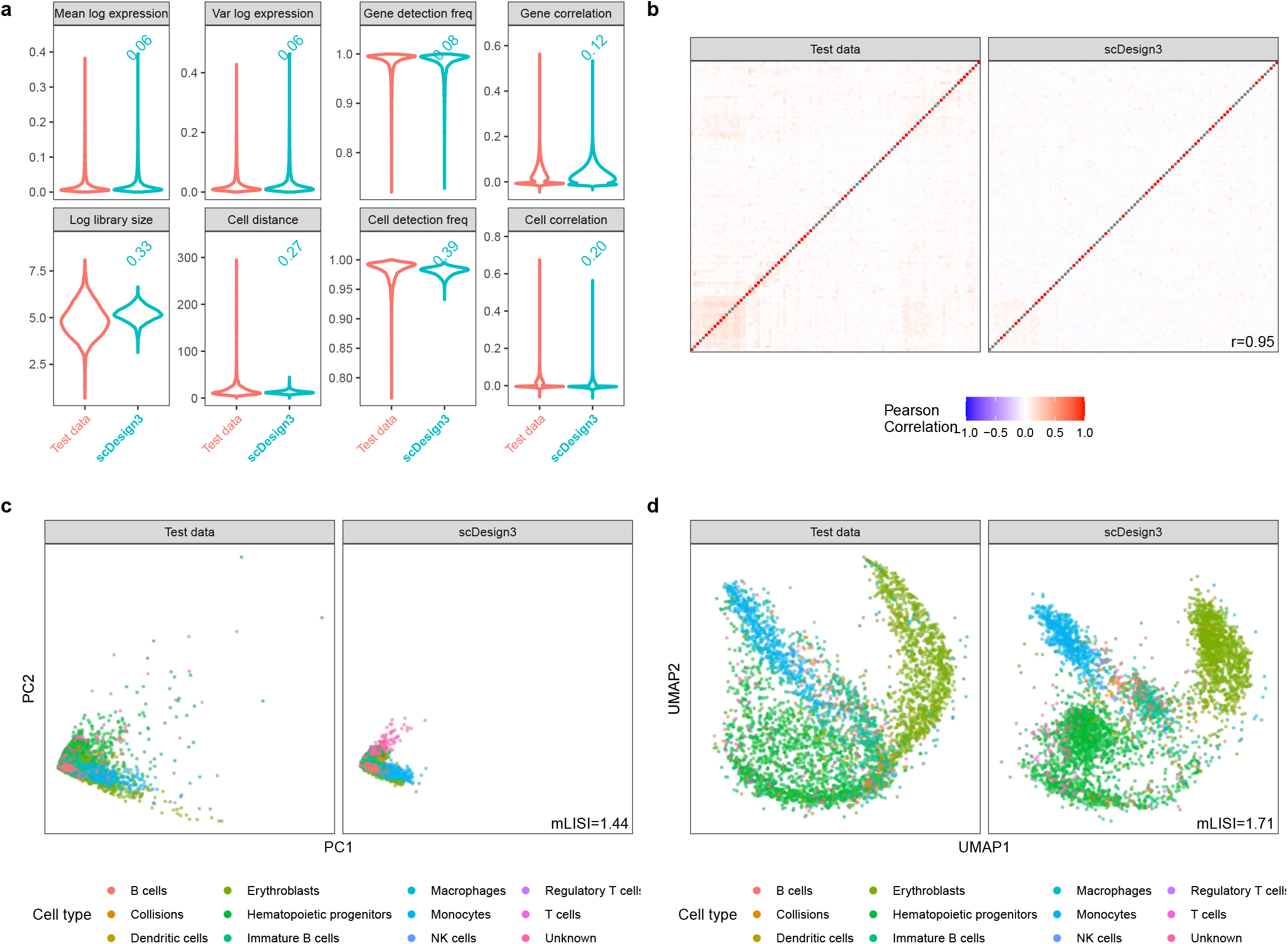
scDesign3 simulates sci-ATAC-seq data (mouse bone marrow). **a**, Distributions of eight summary statistics in the test data and the synthetic data generated by scDesign3 using cell type labels. Each number on top of a violin plot (the distribution of a summary statistic in a synthetic dataset) is the Kolmogorov–Smirnov (KS) distance between the synthetic data distribution (indicated by that violin plot) and the test data distribution. A smaller number indicates better agreement between the synthetic data and the test data in terms of that summary statistic’s distribution.**b**, Heatmaps of the peak-peak correlation matrices in the test data and the synthetic data generated by scDesign3. The Pearson’s correlation coefficient *r* measures the similarity between two correlation matrices, one from the test data and the other from the synthetic data. **c**, PCA visualization (top two PCs) of the test data and the synthetic data generated by scDesign3. The color labels each cell’s cell type. An mLISI value close to 2 means that the synthetic data resemble the test data well in the low-dimensional space. **d**, UMAP visualization of the test data and the synthetic data generated by scDesign3.

**Figure S14:**
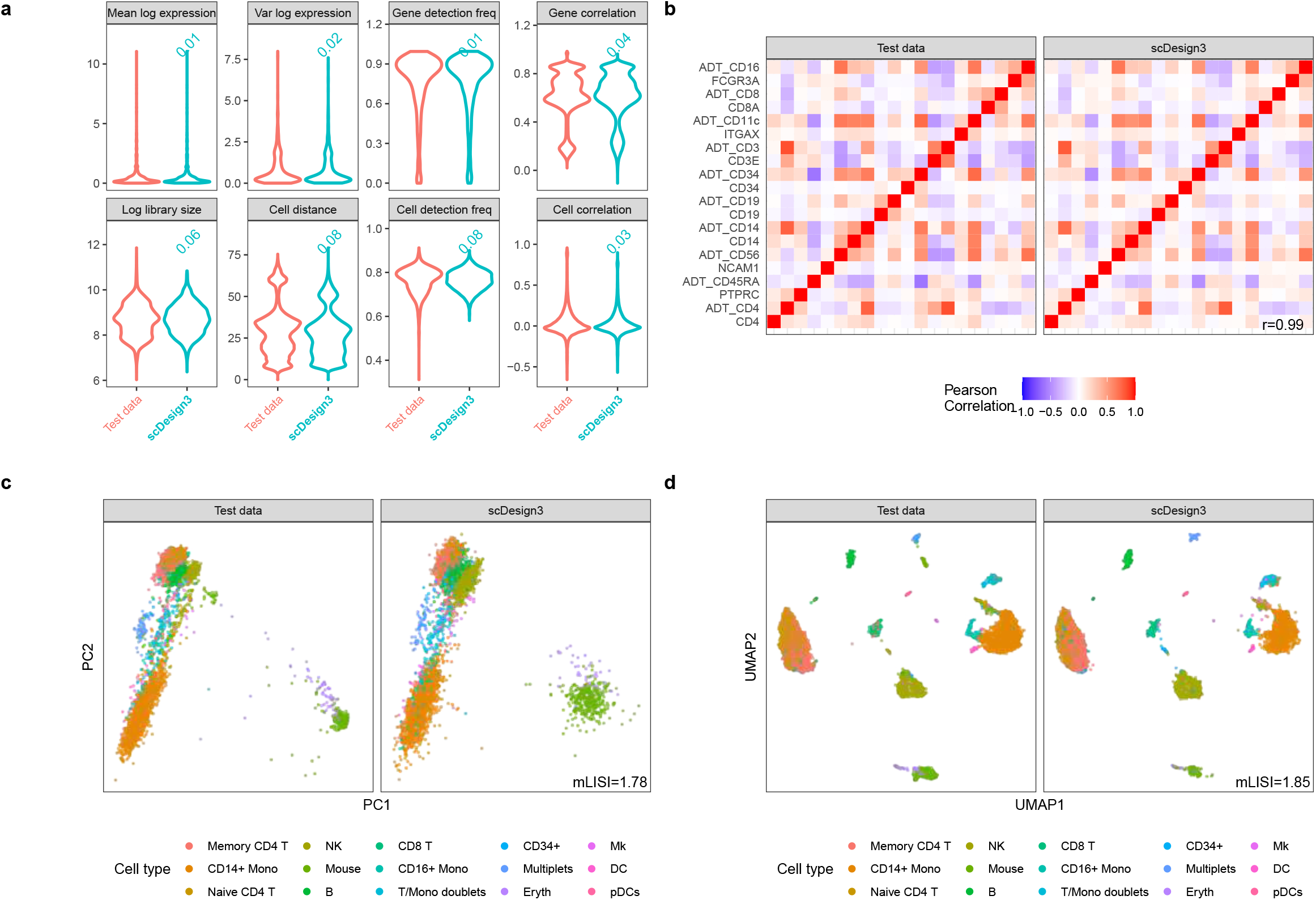
scDesign3 simulates CITE-seq data (human PBMCs). **a**, Distributions of eight summary statistics in the test data and the synthetic data generated by scDesign3. The CITE-seq dataset simultaneously measures each cell’s gene expression and surface protein abundance by Antibody-Derived Tags (ADTs). Each number on top of a violin plot (the distribution of a summary statistic in a synthetic dataset) is the Kolmogorov–Smirnov (KS) distance between the synthetic data distribution (indicated by that violin plot) and the test data distribution. A smaller number indicates better agreement between the synthetic data and the test data in terms of that summary statistic’s distribution. **b**, Heatmaps of the gene and protein correlation matrices (10 proteins with names starting with “ADT” and their corresponding genes) from test data and the synthetic data generated by scDesign3. The Pearson’s correlation coefficient *r* measures the similarity between two correlation matrices, one from the test data and the other from the synthetic data. scDesign3 recapitulates the correlations between the RNA and protein expression levels of the 10 surface proteins. **c**, PCA visualization (top two PCs) of the test data and the synthetic data generated by scDesign3. The color labels each cell’s cell type. An mLISI value close to 2 means that the synthetic data resemble the real data well in the low-dimensional space. **d**, UMAP visualization of the real data and the synthetic data generated by scDesign3.

**Figure S15:**
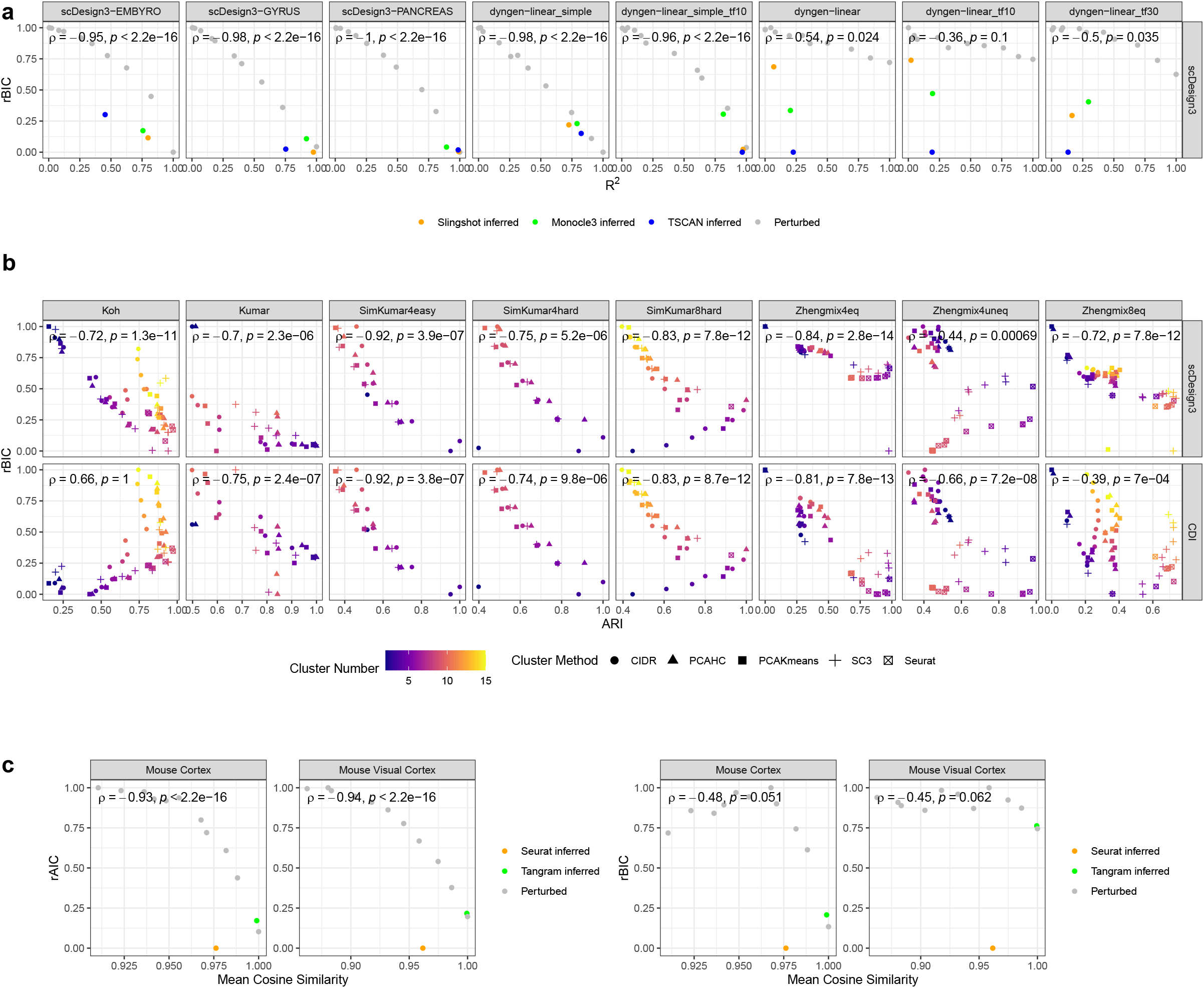
scDesign3 provides an unsupervised quantification of the goodness-of-fit of pseudotime, clusters, and inferred locations. For visual clarity, we plot the relative BIC/AIC (rBIC/rAIC) by re-scaling scDesign3’s marginal BIC/AIC to [0, 1]. **a**, The scDesign3 rBIC (unsupervised) is negatively correlated with the *R*^2^ (supervised) between the perturbed pseudotime plus three inferred pseudotime and the true pseudotime in each of the eight datasets. The true pseudotime is the ground truth used for generating the synthetic data. **b**, Comparison of scDesign3 rBIC and Clustering Deviation Index (CDI) rBIC. The scDesign3 rBIC (unsupervised) negatively correlates with the ARI (supervised). The scDesign3 rBIC has better or similar performance than CDI’s performance on six out of the eight datasets. The color scale shows the number of clusters, and the shapes represent clustering algorithms. **c**, The scDesign3 rAIC (unsupervised) is negatively correlated with the mean cosine similarity (supervised) between the perturbed locations plus two inferred locations, and the true locations in each of the two spatial datasets. The true locations are the ground truth used for generating the semi-synthetic data. Due to the high complexity of spatial patterns, the AIC outperforms BIC since it less penalizes the model complexity.

## 3 Supplementary Tables

**Table S1:** Comparison of scDesign, scDesign2, and scDesign3.

| **Property \ Version** | scDesign | scDesign2 | scDesign3 |
| --- | --- | --- | --- |
| Cell covariate | Cell types<sup>1</sup> | Cell types | Cell types<br>**Cell trajectories**<br>**Spatial locations** |
| Feature type | RNA expr.<sup>2</sup> | RNA expr. | RNA expr.<br>**Chromatin accessibility**<br>**Protein expr.**<br>**DNA methylation** |
| Feature correlation | N/A | Gaussian copula | Gaussian copula<br>**Vine copula** |
| Multi-modality | N/A | N/A | **Multi-omics**<sup>3</sup><br>**Multiple omics**<sup>4</sup> |
| Experimental design | N/A | N/A | **Multiple conditions**<br>**Multiple batches** |
| Feature distribution | Gamma-Normal mixture | Poisson, ZIP<br>NB, ZINB | Poisson, ZIP<br>NB, ZINB<br>**Bernoulli, Normal** |
| Feature mean function | Step function<sup>5</sup> | Step function | Step function,<br>**1D smooth function**<sup>6</sup><br>**2D smooth surface**<sup>7</sup> |
| Model selection | N/A | N/A | **AIC, BIC** |

<sup>1</sup>: scDesign allows cell types to be connected by artificial paths.
<sup>2</sup>: The acronym “expr.” stands for “expression.”
<sup>3</sup>: “Multi-omics” means a cell is measured with multiple modalities.
<sup>4</sup>: “Multiple omics” means a cell is measured with only one modality and more than one modalities are measured on different cells; in this case, scDesign3 requires all cells to be aligned to a common latent space by an integration method.
<sup>5</sup>: Step function is a function of cell type and outputs a constant for each cell type.
<sup>6</sup>: 1D smooth function is a function of cell pseudotime and is modeled by the spline.
<sup>7</sup>: 2D smooth surface is a function of cell spatial location and is modeled by the Gaussian process.

**Table S2:**
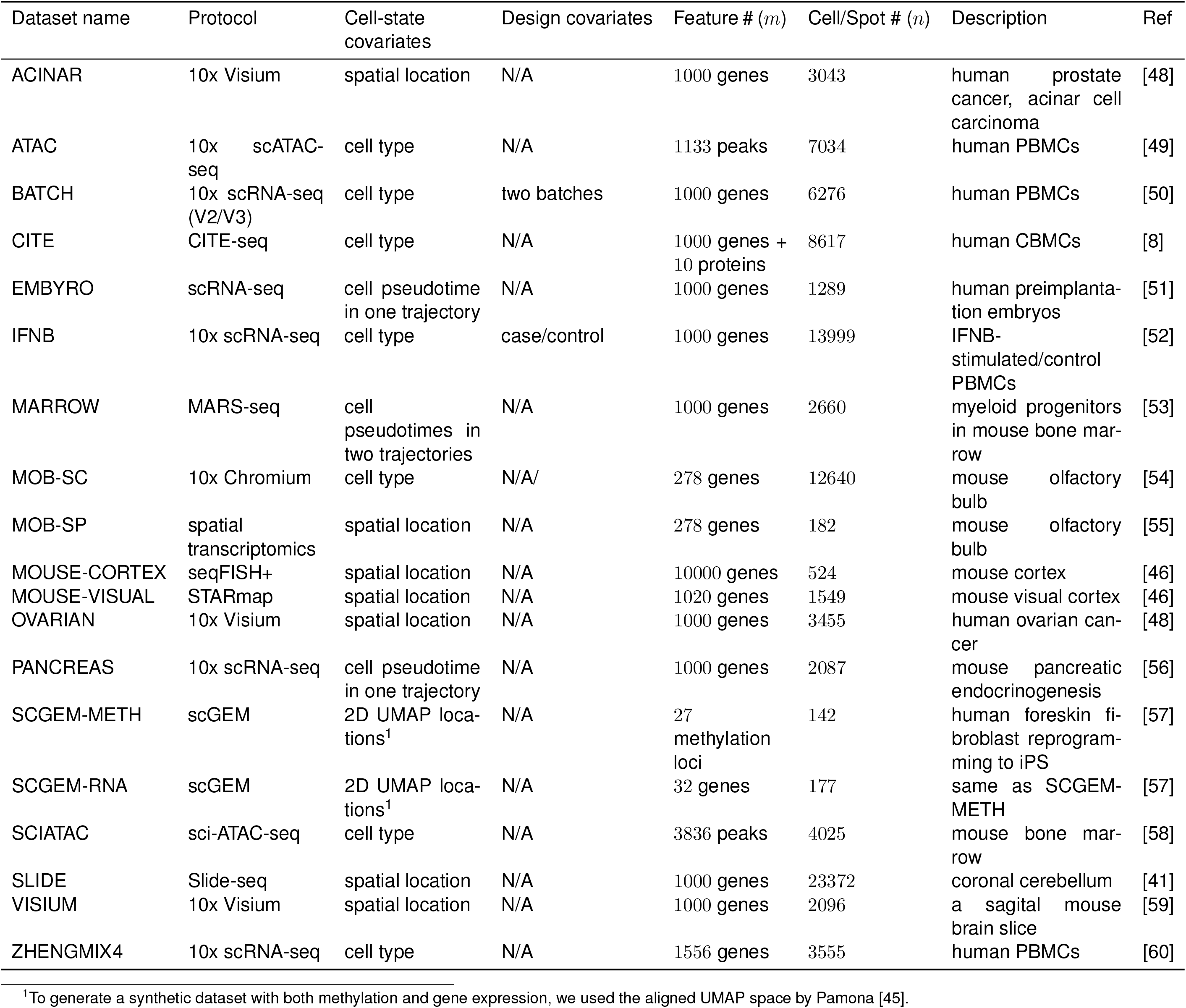
Real datasets used in this study

**Table S3:** Choices of feature *j*’s marginal distribution *F_j_*^1^

| Distribution | Parameters | Probability density function (PDF) or Probability mass function (PMF) | Link function | Applicable data type |
| --- | --- | --- | --- | --- |
| Gaussian | $\mu$: mean<br>$\sigma$: standard deviation | $f(x) = \frac{1}{\sigma\sqrt{2\pi}} e^{-\frac{1}{2}(\frac{x-\mu}{\sigma})^2}; x \in \mathbb{R}$ | $\theta_j(\mu) = \mu$ | Normalized data (e.g., log-transformed count data) |
| Bernoulli | $\mu$: mean | $f(x) = \mu^x(1-\mu)^{1-x}; x \in \{0, 1\}$ | $\theta_j(\mu) = \log \frac{\mu}{1-\mu}$ | Binary data (e.g., DNA methylation) |
| Poisson | $\mu$: mean | $f(x) = \frac{\mu^x e^{-\mu}}{x!}; x \in \{0, 1, 2, \dots\}$ | $\theta(\mu) = \log \mu$ | Count data without over-dispersion |
| Negative Binomial | $\mu$: mean<br>$\sigma$: dispersion | $f(x) = \frac{\Gamma(x+\frac{1}{\sigma})}{\Gamma(\frac{1}{\sigma})\Gamma(x+1)} \left(\frac{1}{1+\sigma\mu}\right)^{\frac{1}{\sigma}} \left(\frac{\sigma\mu}{1+\sigma\mu}\right)^x; x \in \{0, 1, 2, \dots\}$ | $\theta_j(\mu) = \log \mu$ | Count data with over-dispersion (e.g., UMI-based scRNA-seq; spatial transcriptomics; protein abundance) |
| Zero-inflated Poisson | $\mu$: mean<br>$p$: zero-inflation proportion | $f(x) = \begin{cases} p + (1-p)e^{-\mu}; & x = 0 \\ \frac{(1-p)\mu^x e^{-\mu}}{x!}; & x = 1, 2, 3, \dots \end{cases}$ | $\theta_j(\mu) = \log \mu$ | Poisson count data with excess zeros (e.g., scATAC-seq) |
| Zero-inflated Negative Binomial | $\mu$: mean<br>$\sigma$: dispersion<br>$p$: zero-inflation proportion | $f(x) = \begin{cases} p + (1-p)(1+\sigma\mu)^{-\frac{1}{\sigma}}; & x = 0 \\ \frac{(1-p)\Gamma(x+\frac{1}{\sigma})}{\Gamma(\frac{1}{\sigma})\Gamma(x+1)} \left(\frac{1}{1+\sigma\mu}\right)^{\frac{1}{\sigma}} \left(\frac{\sigma\mu}{1+\sigma\mu}\right)^x; & x = 1, 2, 3, \dots \end{cases}$ | $\theta_j(\mu) = \log \mu$ | Negative binomial count data with excess zeros (e.g., full-length non-UMI-based scRNA-seq) |

<sup>1</sup>For notation simplicity, we drop the cell index $i$ and feature index $j$ from all parameters' subscripts.

**Table S4:**
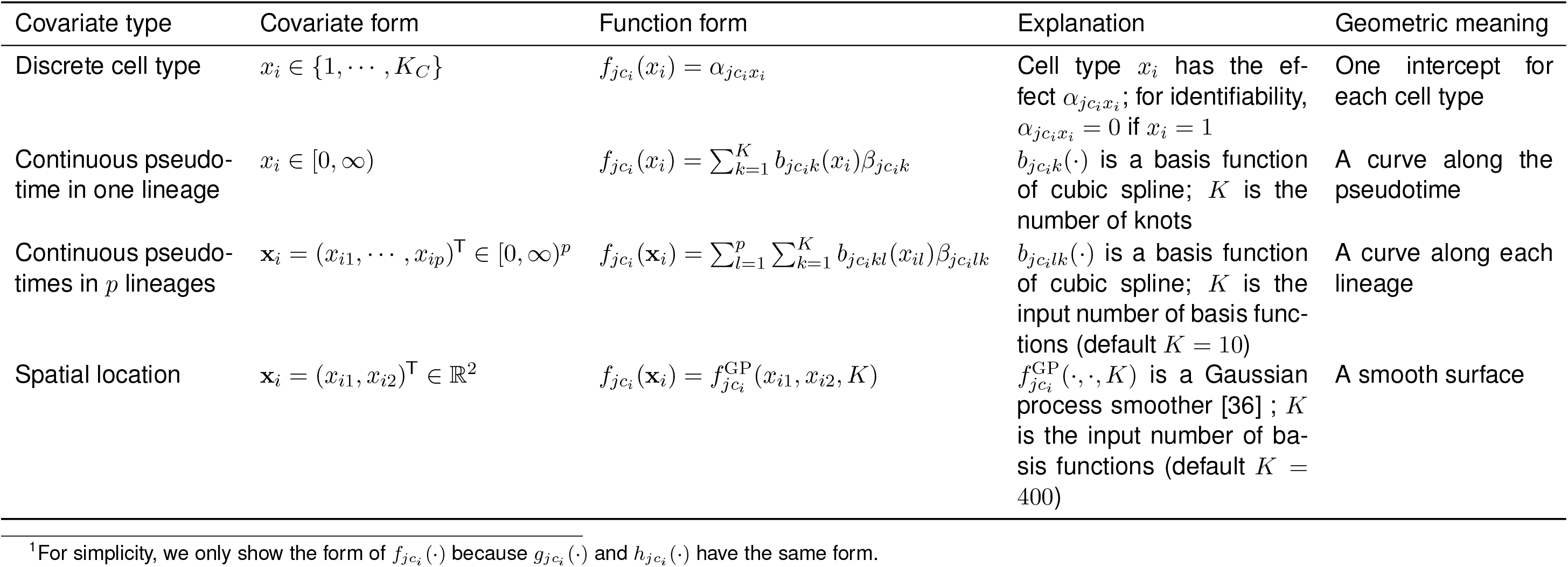
Forms of the functions *f_jc_i__*(·), *g_jc_i__*(·), and *h_jc_i__*(·) of cell-state covariates^1^

**Table S5:**
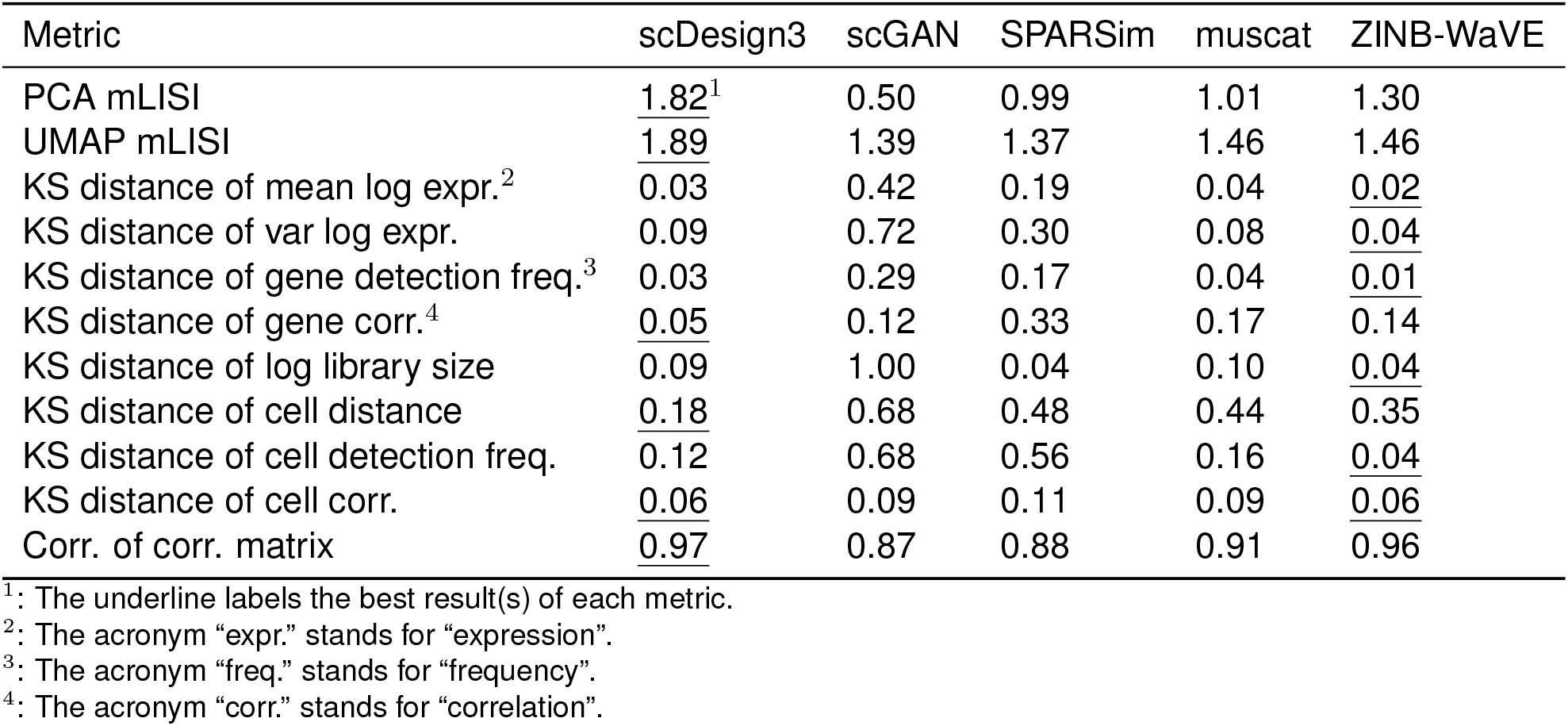
Comparison of scDesign3 and four other simulators (average from PANCREAS, EMBYRO and MARROW)

